# Ultra-sensitive platelet proteome maps the O-glycosylation landscape and charts the response to thrombin dosage

**DOI:** 10.1101/2022.11.01.514776

**Authors:** Callum B. Houlahan, Yvonne Kong, Bede Johnston, Michelle Cielesh, The Huong Chau, Paul R. Coleman, Huilin Hao, Robert S. Haltiwanger, Morten Thaysen-Andersen, Freda H. Passam, Mark Larance

## Abstract

Platelet activation induces the secretion of proteins that promote platelet aggregation and inflammation. However, detailed analysis of the released platelet proteome is hampered by platelets’ tendency to pre-activate during their isolation and a lack of sensitive protocols for low abundance releasate analysis. Here we detail the most sensitive analysis to date of the platelet releasate proteome with the detection of >1,300 proteins. Unbiased scanning for post-translational modifications within releasate proteins highlighted O-glycosylation as being a major component. For the first time, we detected O-fucosylation on previously uncharacterised sites including multimerin-1 (MMRN1), a major alpha granule protein that supports platelet adhesion to collagen and is a carrier for platelet factor V. The N-terminal EMI domain of MMRN1, a key site for protein-protein interaction, was O-fucosylated at a conserved threonine within a new consensus sequence. Our data suggest that Protein O-fucosyltransferase 1 (POFUT1) is responsible for this modification. Secretion of MMRN1 was reduced in cells lacking POFUT1, supporting a key role of O-fucosylation in MMRN1 function. By comparing releasates from resting and thrombin-treated platelets, 202 proteins were found to be significantly released after high-dose thrombin stimulation. Complementary quantification of the platelet lysates identified >3,800 proteins, which confirmed the platelet origin of releasate proteins by anti-correlation analysis. Low-dose thrombin treatment yielded a smaller subset of significantly regulated proteins with fewer secretory pathway enzymes. The comprehensive platelet proteome resource provided here (larancelab.com/platelet-proteome) allows identification of novel regulatory mechanisms for drug targeting to address platelet dysfunction and thrombosis.

**Key Points:**

- High-sensitivity proteome mapping of human platelets identifies O-glycosylation of platelet proteins at key functional sites
- Platelet O-fucosyltransferase POFUT1 regulates the secretion of adhesive protein multimerin-1 (MMRN1)

## INTRODUCTION

Platelets are circulating cells that activate and aggregate after contact with damaged vascular endothelium to promote thrombus formation during haemostasis. It is also well established that platelets play a role in inflammation and vascular repair^1^. Activation, by both chemical and mechanical stimuli, leads to the release of granule contents including soluble proteins, cleaved membrane proteins, and vesicle-bound proteins^2^, which subsequently activates a variety of signalling pathways. Released proteins (i.e. the “releasate”) include those synthesised within the parent megakaryocyte cytoplasm, proteins endocytosed from plasma, as well as proteins synthesised from platelet mRNA^3,4^.

Thrombin activates platelets via the protease-activated receptors PAR1 and PAR4 (F2R and F2RL3)^5-8^. PAR1 mediates platelet activation at low thrombin concentrations (<0.05 U/mL) whereas PAR4 requires a comparatively higher thrombin concentration (>0.1 U/mL) to activate platelets^9,10^. Differential release of pro- and anti-angiogenic factors has been shown with selective stimulation of either PAR1, or PAR4^6^. However, Holten *et al*. identified no qualitative differences upon activation with specific PAR1 and PAR4 agonists^11^. After activation, platelets can release the contents of α-granules, dense granules, tertiary granules, and lysosomes^12^. Platelet α-granules contain adhesive proteins such as fibrinogen (FGA), von Willebrand factor (VWF), multimerin-1 (MMRN1) and fibronectin (FN1), which enable effective thrombus formation. Growth factors (e.g. PDGFA, VEGFC) and chemokines (e.g. CCL5, CXCL3) are also released to regulate the immune response and tissue repair. Platelet lysosomes contain proteases, glycosidases, and acid hydrolases that have bactericidal activity^2^ and may also play a role in receptor cleavage and fibrinolysis^13,14^.

Unbiased mass spectrometry-based proteomics can provide unique insights into cellular protein structure, modifications, and function. Many groups have performed platelet proteomic analysis to understand their protein composition and regulation^2,15,16^. While most proteomic studies have been performed on total platelet lysates or subcellular compartments^17^, the analysis of platelet releasate is much more challenging due to the low concentration of secreted factors. In 2004, Coppinger *et al*. was the first to characterise human platelet releasates after thrombin activation (0.5 U/mL) reporting >300 different proteins using 2D-PAGE^18^. With a novel protein labelling approach a subsequent study identified 124 releasate proteins, following high-dose thrombin (1 U/mL) and collagen (5 µg/mL) stimulation^19^. Using different methods Parsons *et al*. detected 894 released proteins after a similar high-dose thrombin stimulus (1 U/ml)^20^. The differences in these releasate proteomes primarily reflect variance in platelet isolation, platelet stimulation and protein analysis methods. An overlooked aspect of these released proteins is their decoration with post-translational modifications (PTMs) including O-glycosylation, N-glycosylation and proteolytic cleavage, which are important for protein function^21^.

Important questions remain as to the regulation and composition of proteins in healthy human platelets. For example, no study has provided an unbiased view of the PTMs present on platelet releasate proteins. Here, we have addressed these questions using a robust platelet isolation method, coupled to the latest quantitative proteomics and glycomics methodologies for the best sensitivity and accuracy. Subsequent unbiased PTM analysis revealed a wealth of detail in the releasate proteome and highlighted O-glycosylation as a common modification with a large number of unique modification sites identified for the first time. The discovery of O-fucosylation of multimerin-1 (MMRN1) at T216 within its EMI domain provides a novel putative substrate for the platelet glycotransferase enzyme POFUT1. This ultra-sensitive human platelet proteome is shared with the community (larancelab.com/platelet-proteome) to enable future studies in human platelet biology.

## METHODS

Patient blood collection, quality control, and the separation of platelet releasate and lysate for proteomics can be found in the **Supplementary Material**. Human ethics was from the University of Sydney (approval number 2014/244).

### Protein sample preparation for mass-spectrometry-based proteomics

Proteins (5 μg for lysates, 1 μg for releasates, and 70 μg for plasma) were denatured, reduced and alkylated by resuspension in 4% (w/v) sodium deoxycholate (SDC), 10 mM tris-2-carboxyethyl-phosphine (TCEP), 40 mM chloroacetamide, and 100 mM Tris-HCl (pH 8.0), followed by heating to 95°C for 10 min. Samples were then diluted to a final concentration of 1% (w/v) SDC using water and digested for 16 h with trypsin 1:50 (w/w) (Sigma Cat# T6567) at 37°C at 1000 rpm in a Thermomixer-C (Eppendorf). Samples were mixed 1:1 (v:v) with 99% ethyl acetate in 1% (both v/v) TFA and vortexed until all the precipitated SDC was resuspended. StageTips purification of peptides was performed as described^22^. Peptides were reconstituted with 5% (v/v) formic acid in water at ∼0.2 μg/μL and stored at 4°C until liquid chromatography-tandem mass spectrometry (LC-MS/MS) analysis.

### Proteome analysis with LC-MS/MS and data analysis

Peptide samples (0.5 μg) were injected onto a 50 cm x 75 μm C18 (Dr. Maisch, 1.9 μm) fused silica analytical column with a 10 μm pulled tip, coupled online to a nanospray ESI source. Peptides were resolved over a gradient from 5%-35% acetonitrile over 70 min with a flow rate of 300 nL/min. Peptides were ionised by electrospray ionisation at 2.4 kV. MS/MS analysis was performed using a Fusion Lumos tribrid mass spectrometer (ThermoFisher) with higher-energy collisional dissociation. MS/MS spectra were attained in a data-dependent acquisition of the top 20 most abundant ions in each MS1 full scan. RAW data files were analysed using the integrated quantitative proteomics software and search engine MaxQuant^23^ (version 1.6.3.4). A false discovery rate (FDR) of 1% using a target-decoy based strategy was used for protein and peptide identification. The database used for identification contained the Uniprot human database (downloaded 5th of May 2020) alongside the MaxQuant contaminants database. Mass tolerance was set to 4.5 ppm for precursor ions and 20 ppm for fragments. Trypsin was set as the digestion enzyme with a maximum of 2 missed cleavages. Oxidation of Met, deamidation of Asn/Gln, pyro-Glu/Gln, and protein N-terminal acetylation were set as variable modifications. Carbamidomethylation of Cys was set as a fixed modification. The MaxLFQ algorithm was used for label-free quantitation^24^.

### Unbiased (open) PTM search

Data from all high thrombin-treated platelet releasates was combined and searched using Byonic (v3.11.3) ^25^. Initially the search was performed against the whole human proteome database without PTMs. The identified 1,529 proteins were converted into a focused database (**Supplementary File 1**) used for untargeted PTM detection using “wildcard” (open) searches PTMs^26^. An FDR of 1% using a target-decoy based strategy was used for protein and peptide identification. MS1 and MS2 mass tolerance was set to 4 ppm and 20 ppm, respectively. Trypsin was set as the digestion enzyme with a maximum of 2 missed cleavages. Oxidation of Met (common2), deamidation of Asn/Gln (common1), pyro-Glu/Gln(rare1), and protein N-terminal acetylation (rare1) were set as variable modifications. Carbamidomethylation of Cys was set as a fixed modification. Total common max and total rare max were both set to 1. The wildcard search was applied to “unmodified” peptides with a range from -40 to 1000 Da on any amino acid.

### O-glycome profiling

Quantitative O-glycomics analysis of the platelet releasate fractions were performed using an established PGC-LC-MS/MS as previously described^27,28^. See **Supplementary Material** for experimental details and details of the glycomics data analysis including glycan identification and quantitation.

### Statistical analysis

All proteomics data were analysed using R (version 4.03) and plotted using Tableau (version 2020.4). Outliers greater than 1.5 times the inter-quartile range were excluded for some plots to aid visualisation. Fold-changes were calculated based on median values on a per group basis. Statistical significance was determined using a repeated-measures one-way ANOVA for activation (resting vs low/high-dose thrombin), each dose was analysed separately. Outputs were corrected for multiple testing using the Benjamini-Hochberg correction, with significance being set at P<0.05 for an FDR of 5%.

### Data and code availability

Raw MS data have been deposited to the ProteomeXchange Consortium (http://proteomecentral.proteomexhange.org) via the PRIDE partner repository with the dataset identifier PXD036280 (Username, Password: 99h1oIJd). The O-glycan LC-MS/MS raw data files are made publicly available via GlycoPOST^29^, accession number GPST000211.

## RESULTS

### Ultra-sensitive platelet proteome profiling

To examine the platelet proteome with high fidelity we have employed a platelet isolation method that reduces aberrant platelet activation and minimises cellular/plasma contamination (**Figure 1a, 1b and Supplementary Figure 1a**). Based on our thrombin-stimulation dose-response curve for these platelet preparations (**Supplementary Figure 1b**) we defined two doses of thrombin to achieve both low (0.025 U/mL) and high (0.2 U/mL) thrombin stimulation. To confirm platelet activation, we analysed PAC-1 (**Figure 1c**) and P-selectin (**Supplementary Figure 1d**) expression of these cells using flow cytometry. High-dose thrombin yielded >98% P-selectin^+^ and ∼92% PAC-1^+^ cells. In contrast, low-dose thrombin yielded >40% P-selectin^+^ and ∼30% PAC-1^+^ cells.

**Figure 1.**
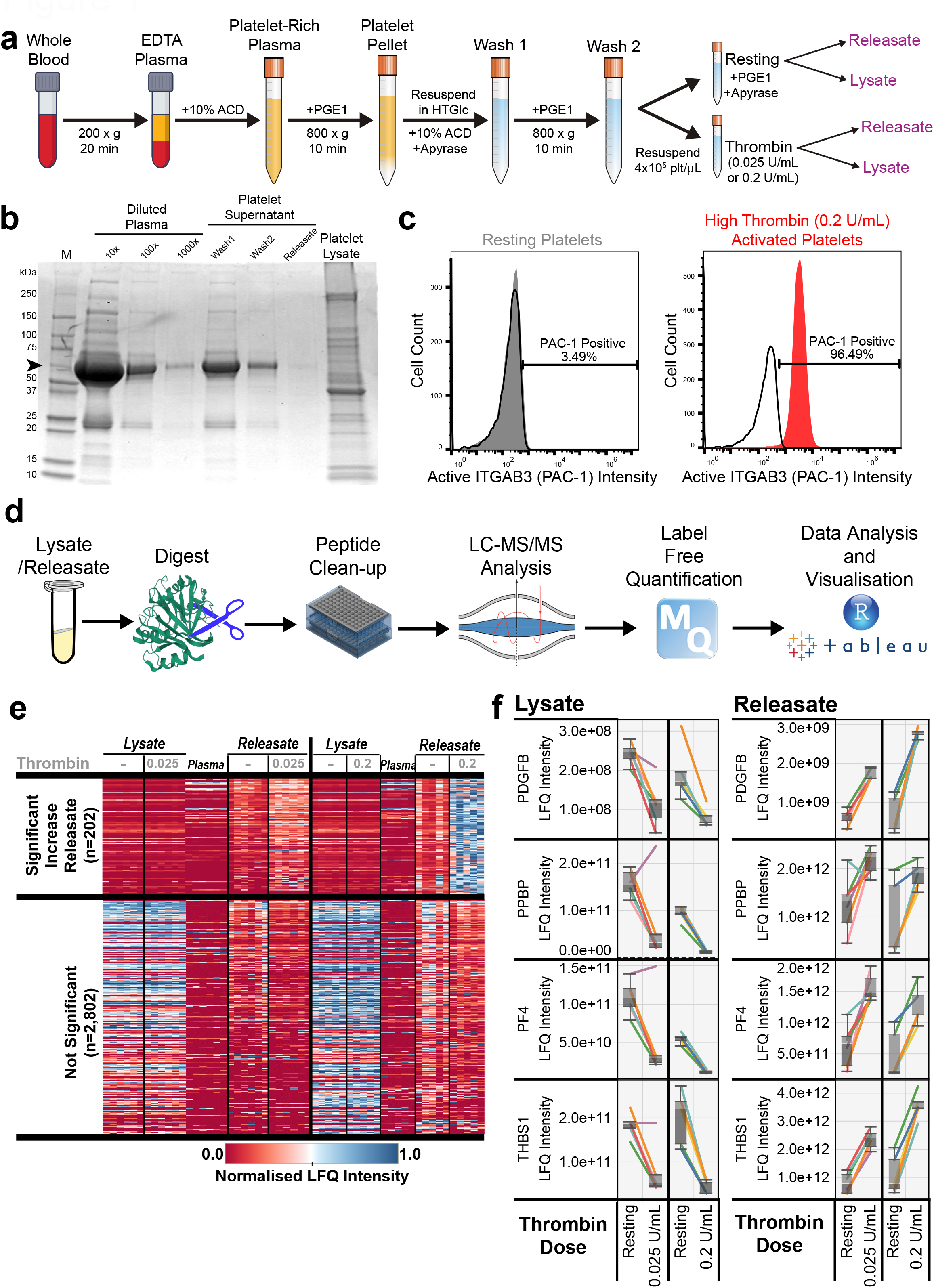
Proteomic analysis of platelet lysates and releasates from healthy donors. (a) Workflow for platelet isolation and stimulation with thrombin. (b) Coomassie stained SDS-PAGE gel to assess for plasma contamination in platelet lysates and releasates. Arrow indicates albumin. (c) Histograms of platelet activation using PAC-1 intensity (x-axis). Resting platelets are shown in grey, platelets stimulated with either low-dose (0.025 U/mL) or high-dose (0.2 U/mL) thrombin shown in red. (d) Workflow for lysate/releasate proteomic analysis. (e) Heatmap showing platelet proteins that were significantly increased in the releasate by high-dose thrombin stimulation, or not significantly regulated, n = 5. (f) Boxplots of proteins known to be secreted by platelets after thrombin activation. Each line represents a single donor. The y-axis shows the label-free quantitation (LFQ) intensity for each protein.

Mass spectrometry-based proteomics was used to examine the platelet lysates and corresponding releasates (**Figure 1d**). Analysis of the platelet releasates identified >1,300 proteins consistently detected across both the high- and low-dose groups (**Supplementary Table 1**). From the platelet lysates >3,000 proteins were consistently detected (**Supplementary Table 1**). To visualise the thrombin-induced proteome changes, we plotted a heat map of the normalised label-free quantitation (LFQ) intensity for all proteins detected (**Figure 1e**). Proteins known to be released upon thrombin activation, showed the expected responses after thrombin stimulation (**Figure 1f**).

### Unbiased PTM analysis of releasate proteins

PTMs are known to play important roles in the function of platelet releasate proteins. For example, the O-glycosylation of thrombospondin (THBS1) has been proposed to mediate protein-protein interactions and protein folding^30^. To investigate modifications to proteins in the releasate proteome, we employed an open-search strategy^26^. Peptide-level MS/MS data from all thrombin-treated platelet releasates were combined and searched for modifications ranging in mass from -40 to +1000 Da, representing a wide range of potential PTMs (**Supplementary Table 2)**. As expected, we detected chemical modifications of <100 Da arising during sample preparation, such as oxidation (+16 Da) (**Figure 2a**).

**Figure 2.**
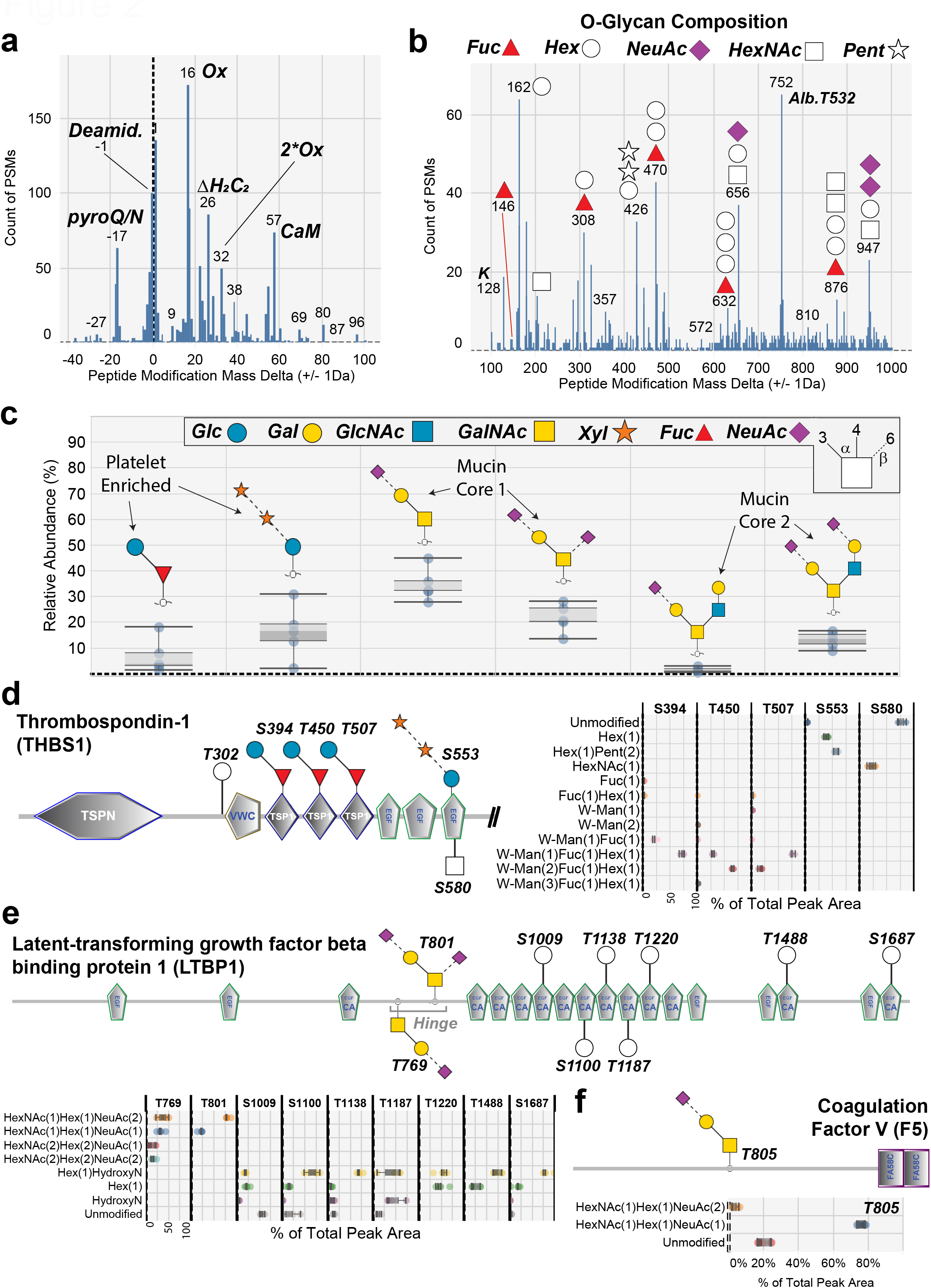
Unbiased detection of protein modifications including O-glycosylation of platelet releasates. Histograms of the open-search analysis of high-dose thrombin stimulated platelet releasates showing the number of peptide spectral matches (PSMs) (y-axis) across all platelet samples identified as containing the indicated mass adducts (x-axis). Modifications were plotted separately for common low-mass modifications (a) and higher mass modifications (b). Delta masses corresponding to O-glycan masses are indicated by the glycan symbols. (c) Quantitative O-glycomics analysis of O-glycans detached by β-elimination from platelet releasate proteins as described in Supplementary Methods (n=5). Bond linkage types are indicated in the legend. (d) Boxplots showing quantitative analysis of thrombospondin-1 O-glycosylation at identified sites, (n=5). W-Man is C-mannose modified tryptophan on the same peptide as the indicated Ser/Thr. (e) Boxplots showing quantitative analysis of latent-transforming growth factor beta-binding protein 1 O-glycosylation at identified sites, (n=5). HydroxyN is hydroxylated asparagine on the same peptide as the indicated Ser/Thr. (F) Boxplots showing quantitative analysis of coagulation factor V O-glycosylation at identified sites, (n=5).

Unexpectedly, plotting peptide modifications >100 Da revealed ∼20 prominent peaks, whose masses corresponded with known O-glycan compositions and were associated with Ser/Thr residues^31^ (**Figure 2b**). Of these, four modifications contained fucose in addition to hexose, pentose, and HexNAc residues^32^. Two modifications had compositions consistent with mucin-type sialo-O-glycans. Peptides modified by fucose alone (146 Da) were identified, alongside many peptides modified by a single hexose (162 Da). In total 36 proteins were identified as being O-glycosylated including a small group of proteins modified at multiple sites including THBS1, latent-transforming growth factor beta-binding protein 1 (LTBP1), fibrinogens (FGA/FGB) and platelet basic protein (PPBP). The discovery that these proteins carry O-glycans is important as this small protein subset constitutes >50% of the protein content released by platelets (**Supplementary Table 2**).

The corresponding O-glycan profile was established by quantitative O-glycome analysis, where all O-glycans are detached through β-elimination from already de-*N*-glycosylated proteins and profiled by LC-MS/MS^33^ (**Figure 2c**). This O-glycome profiling method provides both the fine structures and relative abundances of O-glycans. O-glycans with 11 different structures covering 9 different compositions were identified as the main components attached to platelet releasate proteins (**Supplementary File 2**). These glycans quantitatively agreed with our open search analysis of the proteomics data, as we observed a high abundance of O-fucose structures extended by glucose, (xylose)_2_-glucose conjugates, and mucin-type core 1 and 2 sialo-O-glycans (**Supplementary Table 3**).

Of the O-glycosylated sites detected on releasate proteins many were not previously observed in human samples including platelets. To validate these PTMs we performed electron-transfer/higher-energy collision dissociation (EThcD) fragmentation analysis on a separate cohort of platelet samples (**Supplementary Table 4**). EThcD fragmentation-based MS/MS analysis is known to preserve labile modifications such as O-glycans for more confident identification^34^. Among the many releasate proteins confirmed to be O-glycosylated, four modified proteins were the most novel and were annotated in the context of their known domains^35^. First, we observed that ∼53% of S553 of THBS1 had a (pentose)_2_-hexose modification consistent with a (xylose)_2_-glucose conjugate (**Figure 2d**). Second, we observed two sialylated mucin-type O-glycans on LTBP1 at T769 and T801 within the proteolytically sensitive hinge domain, which targets the protein to the extracellular matrix and is needed for TGF-beta release^36^ (**Figure 2e**). Third, coagulation factor V (F5) carried mostly a core 1 mucin O-glycan (>75% sialyl T) in position T805 while ∼20% of the site was unmodified (**Figure 2f**). Lastly, we observed >95% O-fucosylation of multimerin-1 (MMRN1) at T216, which is outside the EGF-like and TSR domains known to contain this modification^37^.

### Quantification of releasate proteins

Overall, we identified 202 proteins with significantly increased total abundance in the high-dose thrombin releasates, whereas only 63 were significantly increased by the low-dose thrombin (**Supplementary Table 5**). To delineate the functional groups of the proteins increased in high-dose thrombin releasates, we categorised proteins into 10 distinct gene ontology groups based on either biological process, or cellular component (**Figure 3**). As expected, proteins known to reside in the platelet alpha granules underwent the largest fold-change increase and were also the most abundant releasate constituents. This was closely followed by proteins involved in cellular signalling such as chemokines and growth factors (e.g. C-X-C motif chemokine 3 - CXCL3, C-C motif chemokine 5 - CCL5). In this signalling group, several proteins not previously described to be released from platelets were detected, including granulin (GRN) and midkine (MDK), with granulin having the largest fold-change and final protein abundance. The group showing the largest average fold-change were from the lysosome^38^. Unexpectedly, many proteins from the Golgi and endoplasmic reticulum were also observed in the releasate, a large proportion of which are involved in glycan processing/metabolism^39^. Of the 133 proteins having accurate measurements of lysate fold-change, only 44 decreased in lysate abundance after high thrombin by >2-fold (**Figure 4a**,**b** and **Supplementary Table 5**). Several lysosomal proteins that were enriched in releasates (**Figure 4c**), such as N4-beta-N-acetylglucosaminyl-L-asparaginase (AGA) showed a 32-fold increase in platelet releasates after thrombin, but only <2-fold decrease in their lysate abundance (**Figure 4d**). This suggests a large pool of these proteins likely remains within the cell and that only a small portion is secreted after thrombin treatment. Lastly, proteins that mediate cell-cell contacts and comprise the extracellular matrix were also released and included proteins such as glycoprotein V (GP5)^40^ and nidogen 1 (NID1), a basement membrane component.

**Figure 3.**
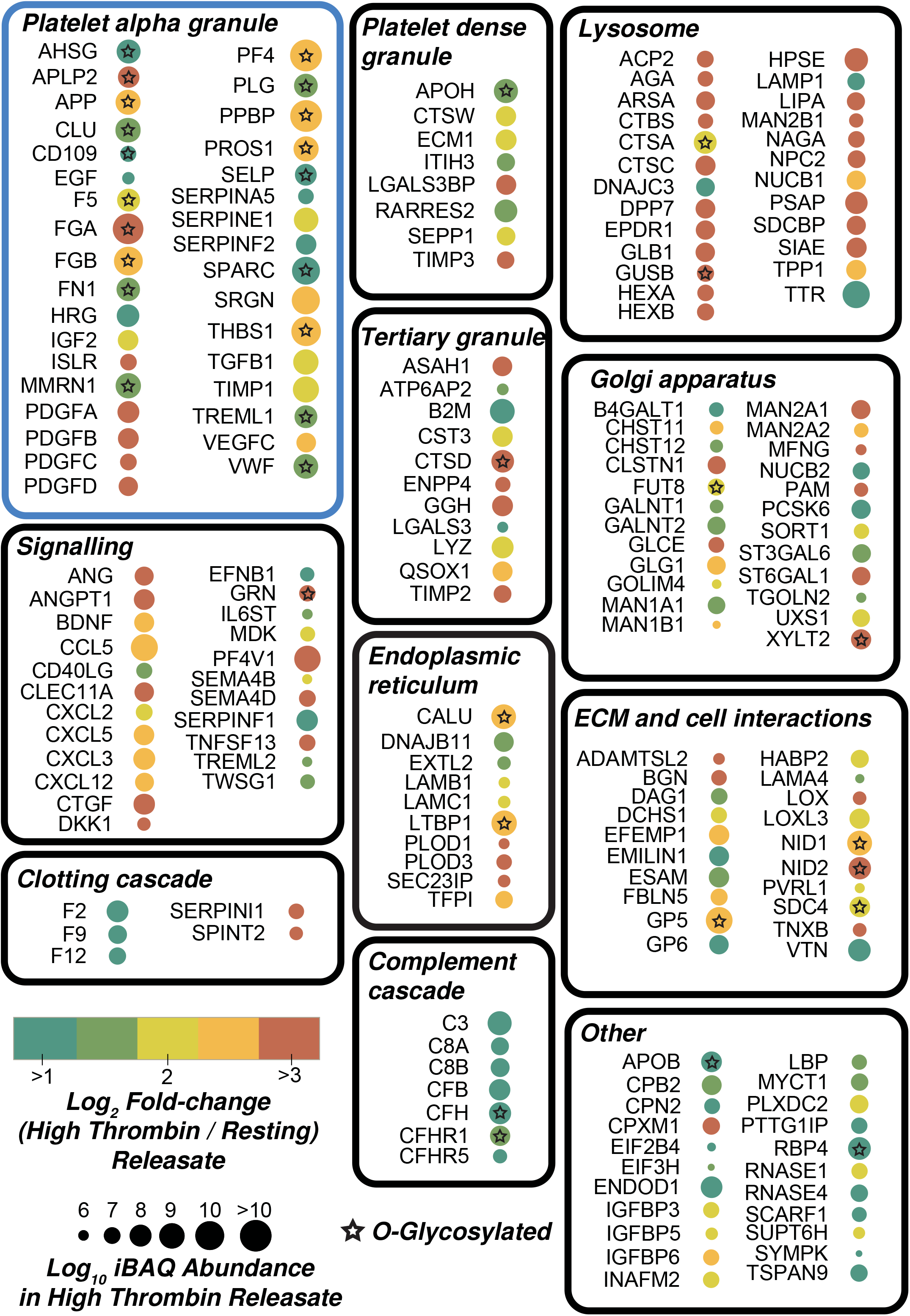
Functional grouping and response of platelet proteins significantly increased in the releasate after high-dose thrombin stimulation. Each protein is represented by a circle annotated with the gene name, and the circle colour represents the log_2_ fold-change (thrombin stimulated/resting) in the releasate (i.e. the degree of change). The circle size represents the log_10_ iBAQ abundance of each protein in the high-dose thrombin stimulated releasate (i.e the proportion of the protein relative to the total proteins). Stars indicate releasate proteins that were identified to be O-glycosylated.

**Figure 4.**
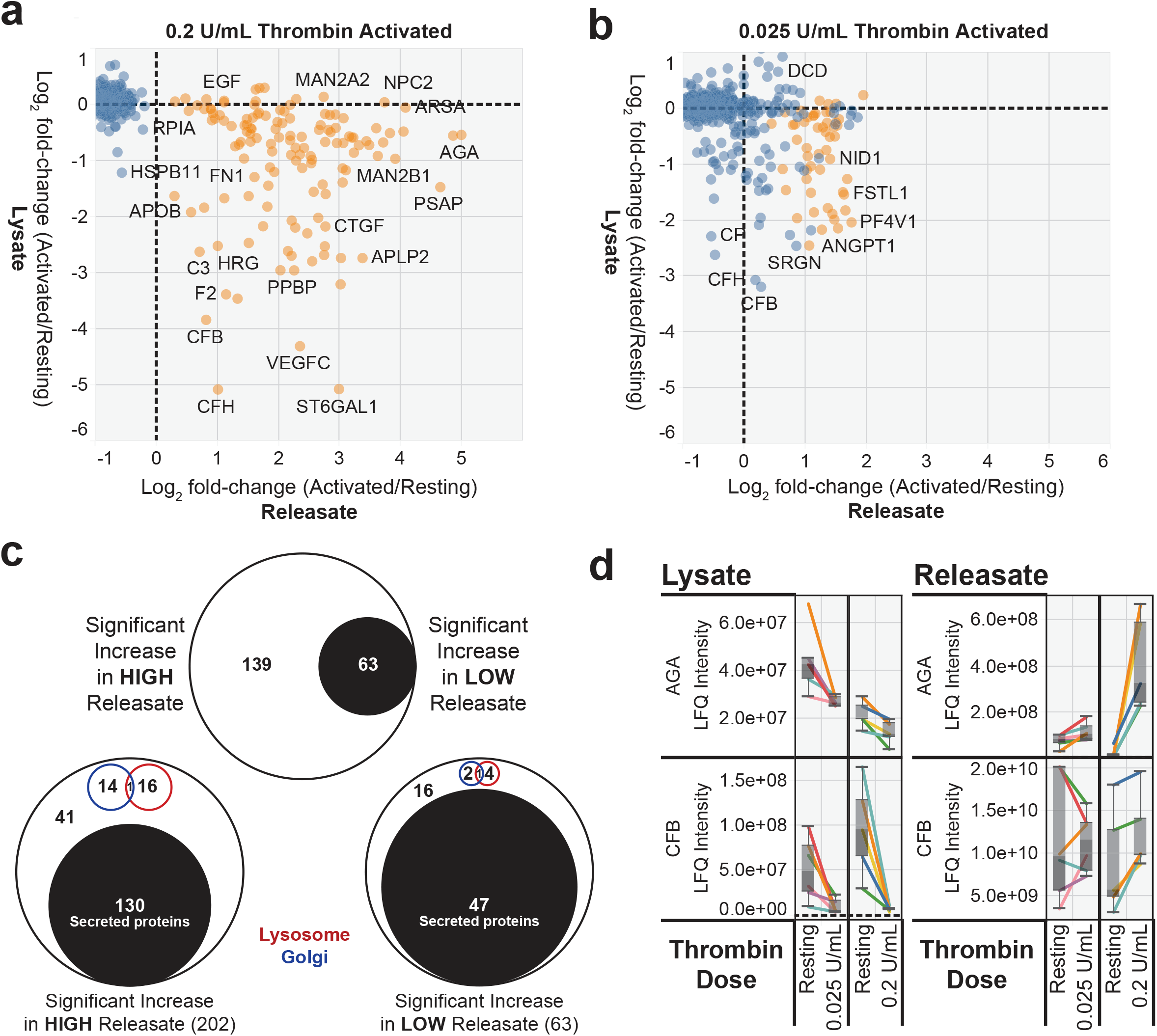
Comparison of platelet lysate and releasate proteomes for confident detection of released proteins from thrombin-activated platelets. Scatterplot of proteins significantly regulated in the high-dose (a) and low-dose (b) thrombin stimulated releasate. Significantly regulated proteins are shown in orange, non-significant proteins are shown in blue. The log_2_ fold-change (thrombin stimulated/resting) for both the each corresponding releasate (x-axis) and the corresponding lysate (y-axis) was used for plotting. (c) Venn diagrams indicating the overlap between significantly increased releasate proteins in either the low-dose or high-dose thrombin groups, *top*. Analysis of the proportion of significantly increased releasate proteins that are annotated by Uniprot as secreted, Golgi-associated, lysosome-associated, or other. (d) Boxplots of proteins significantly regulated in platelet releasates-only (AGA) or lysates-only (CFB) after thrombin activation. Each line represents a single donor. The y-axis shows the label-free quantitation (LFQ) intensity for each protein.

### Characterisation of multimerin-1 fucosylation within the EMI domain

One of the most abundant proteins released by activated platelets was MMRN1, a large ∼150 kDa protein that initially forms stable homotrimers, which are hypothesised to interact through EMI domain and C1q domain binding to form elongated multimers of up to many MDa in size^41^. We observed a novel fucosylation of MMRN1 at T216, which is within the N-terminal EMI domain of the protein. The O-fucosylation on T216 of MMRN1 was confirmed by EThcD analysis as was the predicted O-fucosylation of the C-terminal EGF-like domain at T1055 (**Figure 5a and Supplementary Table 4**). The stoichiometry of T216 fucosylation is very high, as the fucosylated peptide was >10-fold more intense compared to the non-glycosylated form (**Supplementary Table 2**). The modified T216 and its surrounding sequence are highly conserved in vertebrates (**Figure 5b**) and the EMI domain has 6 conserved cysteines, which likely form disulfide bonds^42^. To examine the predicted 3D-structure of the MMRN1 EMI domain we used the RoseTTAfold program^43^, which predicted the correct pairing of cysteine residues^44^ (**Figure 5c**). Mapping the modified threonine residue onto this structure illustrates the small size of the EMI domain relative to the fucose modification (*pink circle*, **Figure 5c**). The EMI domain is hypothesised to contain 2 sub-domains, where one is similar to the C-terminal region of the EGF-like domain^42^. EGF-like domains are known to be O-fucosylated by the ER resident enzyme GDP-fucose protein O-fucosyltransferase 1 (POFUT1)^45^. O-fucosylation of EGF-like domains by POFUT1 occurs in a specific motif (C^2^XXXX[S/T]C^3^) bracketed by the second and third conserved cysteine residues^37,46^. In contrast, the related enzyme POFUT2 adds O-fucose to TSRs containing CXX[S/T]C motifs such as THBS1 as shown in **Figure 2d**^46^. Sequence similarities were observed after alignment of the MMRN1 EMI-domain with EGF-like domains that are known to be O-fucosylated by POFUT1 (**Figure 5d**). This showed T216 of MMRN1 is in a region of the EMI-domain not conserved with EGF-like domains, but displays a similar sequence motif that is only missing the C-terminal cysteine residue and would be compatible with transfer of fucose^47^. The region of MMRN1’s EMI domain that is homologous to the EGF-like domain fucosylation site does not match the POFUT1 modification motif, as one extra amino acid has been inserted and this is known to be incompatible with fucosylation^47^. Therefore, this modification of MMRN1 at T216 likely represents a new recognition motif for POFUT1 within a new domain type that is missing the C-terminal cysteine residue.

**Figure 5.**
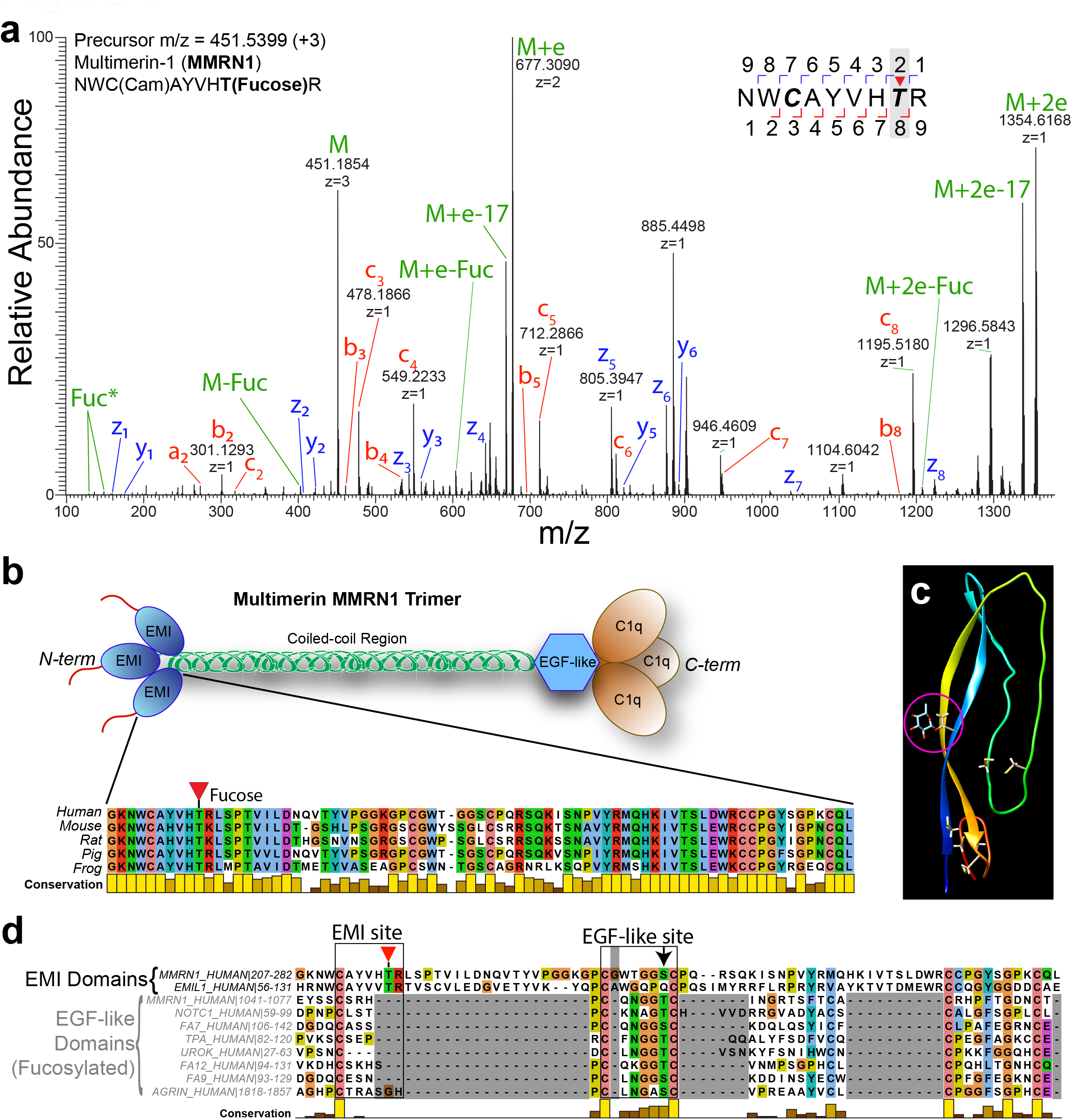
Identification of a novel O-fucosylation site on platelet multimerin 1. (a) Electron transfer higher collision energy dissociation (EThcD) mass spectrum of the novel O-fucosylation site at T216 in MMRN1. The intact precursor ions and associated neutral losses are in green, the z and c fragment ion series are shown in blue and red, respectively. (b) Cartoon of the trimeric MMRN1 structure with key domains shown. *Inset*, protein sequence alignment of the MMRN1 EMI domain across diverse species. (c) Structural prediction of the EMI domain from RoseTTafold with cysteine residues shown as ball and stick and the fucosylated threonine residue highlighted inside a pink circle. Cysteine residues are highlighted by showing the side-chain atoms (d) Protein sequence alignment of human EMI domains and EGF-like domains that are known to be O-fucosylated by POFUT1. The EMI site is highlighted with a red triangle indicating the site of O-fucosylation. The EGF-like site is highlighted with the black arrow indicating the modified residue in EGF-like domains.

Using our proteomics dataset, POFUT1, but not the related enzyme POFUT2, was detected in the platelet lysates and underwent no significant change in abundance after thrombin stimulation (**Figure 6a**). To determine the extent of possible POFUT1 targets, we aligned all human EMI domains (**Figure 6b**). This showed EMILIN1, multimerin-2 (MMRN2), EMI domain-containing protein 1 (EMID1), and collagen alpha-1(XXVI) chain (COL26A1) had the highest similarity and are also potential O-fucosylation targets of POFUT1. Of these, only EMILIN1 was detected in our platelet releasates. We observed O-fucosylation at the conserved position T65 within the EMI-domain of EMILIN1, which is surrounded by the same motif as seen for MMRN1 (CXXXX[S/T]X) (**Supplementary Table 2**). Together, these findings suggest EMI-domains likely are a previously uncharacterised target of O-fucosylation by POFUT1 and that these modifications may play a key role in the function of MMRN1. A known function of O-fucosylation is to assist protein folding, however exogenous expression in HEK293 cells of myc-tagged MMRN1 that was either wildtype, or a T216A mutant, showed no difference in lysate MMRN1 protein abundance (**Figure 6c**). To examine the role of POFUT1 in MMRN1 secretion we expressed myc-tagged wild type MMRN1 in HEK293T cells that were either wild type, *POFUT1*-null mutants, or *POFUT2*-null mutants. Fucosylated wildtype MMRN1 could be detected in the tissue culture media from wild type and POFUT2 null cell lines, but was not detected in the POFUT1 null cells (**Figure 6d**).

**Figure 6.**
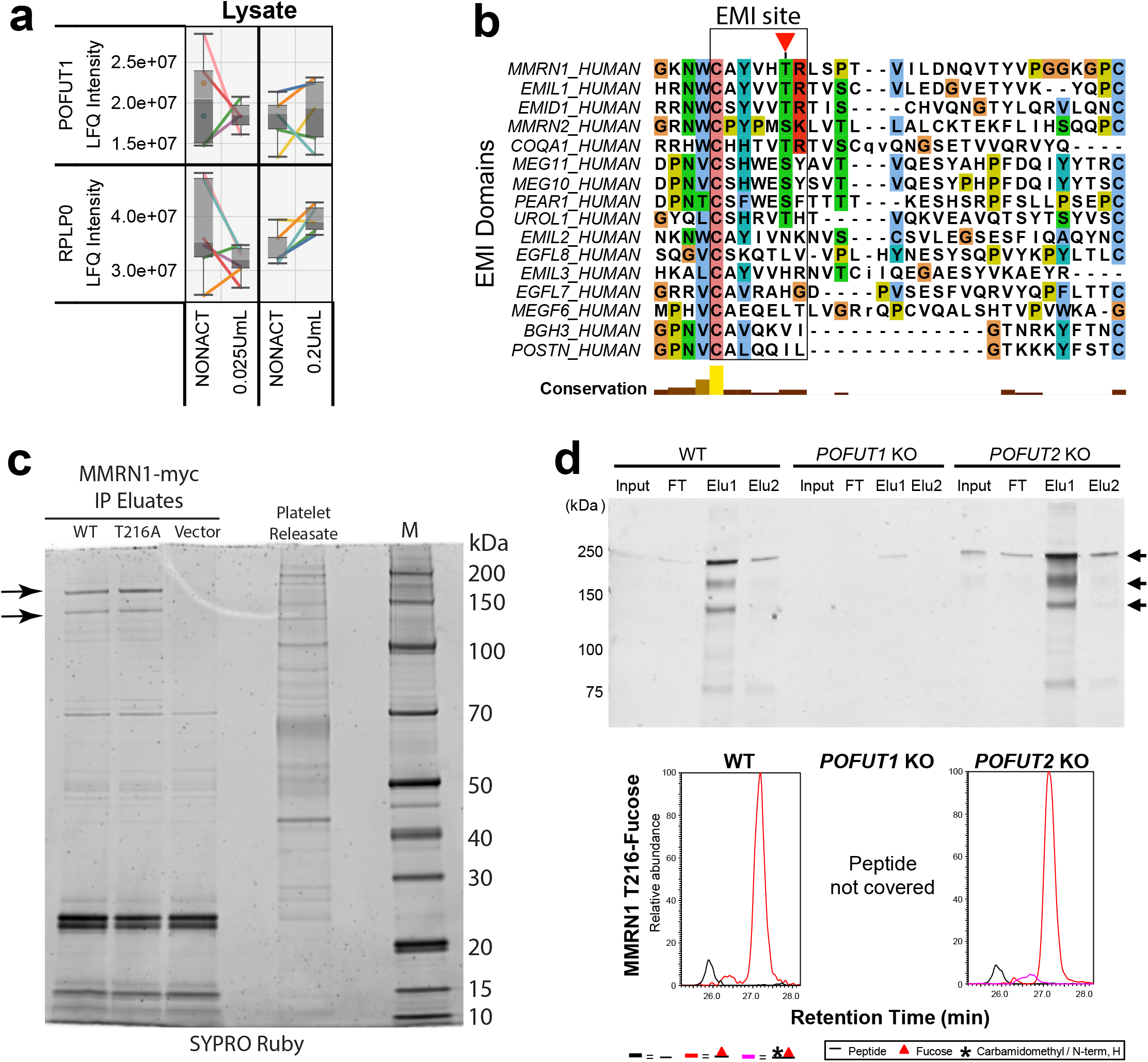
Loss of POFUT1 affects multimerin-1 secretion. (a) Boxplots of POFUT1 and the negative control RPLP0 (an abundant ribosomal protein) in platelet lysates before and after thrombin activation. Each line represents a single donor. The y-axis shows the label-free quantitation (LFQ) intensity for each protein. (b) Protein sequence alignment of EMI domains across a wide range of human proteins. The EMI site is highlighted with a red triangle indicating the site of O-fucosylation. (c) SYPRO Ruby stained SDS-PAGE gel showing the immunoprecipitated (IP) myc-tagged MMRN1 from HEK-293 cells. Cells express either empty vector, wild type MMRN1, or a T216A mutant form of MMRN1. Black arrows indicate MMRN1. (d) *Top*, western blot for myc-tagged wild type MMRN1 that was purified by Ni-NTA agarose from culture supernatants of transfected HEK-293 cells. HEK-293 cells were either wild type (WT), POFUT1 knock-out (KO), or POFUT2 KO. *Bottom*, extracted ion chromatograms from LC-MS/MS analysis using targeted mass spectrometry acquisition for the unmodified T216-containing peptide (black lines), or the O-fucosylated form (red lines).

## DISCUSSION

In this study, we comprehensively mapped thrombin-induced platelet responses by combining high-quality platelet preparations and ultra-sensitive mass spectrometry-based proteomics and glycomics. This enabled the identification of new factors being released by platelets and observation of novel PTMs on platelet proteins. Our analysis generated three key findings. First, platelet releasate proteins were enriched for a wide range of O-glycan modifications and we quantified many novel modification sites and structures. Second, the abundant platelet releasate protein multimerin-1 (MMRN1) was determined to be O-fucosylated on a novel site (T216) within its EMI-domain, and we show that fucosylation by POFUT1 is needed for its secretion. We propose EMI domains form the second-known substrate of platelet POFUT1 that indicates a much more widespread functional role for this enzyme in platelets. Lastly, large differences in platelet secretion response were observed between low-dose and high-dose thrombin, with strong stimuli triggering increased secretion of lysosomal luminal enzymes. Collectively, these data provide a comprehensive proteome-wide analysis of platelet responses to thrombin activation, which may contribute to disease states through new functions in haemostasis. This proteomic resource is provided as a free web-based interactive visualization for the research community (<u>larancelab.com/platelet-proteome</u>).

O-glycosylation is known to play an important regulatory role in cellular function, however, the majority of platelet studies have focused on the role of N-glycans and O-sialic acids in platelet production and clearance^48,49^. Defective O-glycosylation of platelet proteins results in a bleeding phenotype^50^. The recent progress in mass spectrometry technology has enabled the site-specific detection of labile O-glycan modifications with high sensitivity and confidence in human samples^34,51^. Previous studies have used enrichment regimes to facilitate the detection of platelet O-glycosylation^52^. However, these regimes often only allow the identification of specific glycan classes (e.g. O-GalNAc glycans) and do not enable identification of some platelet-enriched O-fucosylation, or O-glucosylation. Here, we demonstrate that many of the released platelet proteins carry diverse O-glycan structures. This included high abundance of O-fucosylation and O-glucosylation and their extended forms, alongside the more broadly expressed mucin-type glycans. The high abundance of O-fucosylated proteins can be explained through domain enrichment analysis of the >200 platelet releasate proteins we detected using STRING-db^53^, where 19 proteins (e.g. THBS1, MMRN1, LTBP1, NID1) contained either EGF-like domains, or TSRs, which are the only previously known consensus sites for O-fucosylation. This study has now demonstrated for the first time that many of those proteins/sites that were only predicted to be modified, such as the EGF-like domains in several proteins^37^, are indeed O-fucosylated at high stoichiometry in human platelets and are likely to contribute to platelet function.

We have discovered that the abundant platelet releasate protein MMRN1 is constitutively O-fucosylated at T216 within its EMI domain. MMRN1 is known to have several functions including binding/release of coagulation factor V, thrombus formation^54^ and interaction with collagen via an N-terminal RGD motif^55^. We show that T216 was stoichiometrically fucosylated within the context of a potentially altered POFUT1 O-fucosylation motif (C^1^-X-X-X-X-T-X) derived from its known modification context in EGF-like domains^47^. We show that POFUT2 is not involved in O-fucosylation of T216, so it is likely that POFUT1 is responsible. Nonetheless, confirmation of this awaits direct demonstration that POFUT1 can modify this site. Alternatively, a novel POFUT may exist that modified T216. Analysis of MMRN1 secretion in a POFUT1 null cell line showed a complete inhibition of secretion suggesting that O-fucosylation plays a role in MMRN1 secretion. While O-fucosylation was not required for protein expression in HEK293 cells, it may be required for correct folding of the tertiary or quaternary structure of MMRN1, which exists natively as a trimer^41^. Given the proposed role of the N-terminal EMI domain to interact with the C-terminal C1q domain and enable MMRN1 multimerization^42^, we propose that O-fucosylation is required for this interaction and is important for MMRN1 secretion and possibly multimerization efficiency.

High sensitivity mass spectrometry-based proteomics of the platelet releasate has enabled us to identify several novel factors released from platelets in response to high-dose thrombin. These include two small, secreted proteins granulin (GRN) and midkine (MDK). Given the known links between lysosomal function and granulin activity^56,57^, we hypothesise that granulin plays a key role in the formation of the platelet lysosomes. Midkine signalling through a range of receptors is known to control inflammatory processes including the activation of neutrophils^58^. Therefore, we hypothesise that the release of the pro-inflammatory MDK cytokine by platelets contributes to the neutrophil recruitment at the site of tissue damage and, in turn, enhances the innate immune defence.

This work comprehensively catalogues the proteome-wide response of human platelets to thrombin activation. Platelets experience a range of thrombin levels *in vivo*; the thrombin concentrations during coagulation are estimated to range from 1 nM (0.1 U/mL) to over 500 nM^59^. Different thrombin concentrations will also affect the platelet interacting environment including the composition of fibrin strands^60^. Alterations in the platelet releasate content in response to different thrombin concentrations, may differentially modulate platelet functions including immune, inflammatory, angiogenic and tissue remodelling responses^61^.

We show that high-dose (0.2 U/mL) thrombin can trigger the release of many proteins associated with the secretory pathway including the ER, Golgi and lysosome. Our comparison of platelet lysate and releasate fold-changes after thrombin treatment, showed that only a small pool of these secretory pathway proteins was released from platelets. Furthermore, several low abundance proteins detected in thrombin-stimulated releasates such as tumor necrosis factor ligand superfamily member 13 (TNFSF13) and calsyntenin-1 (CLSTN1), could not be reliably detected in the corresponding lysates^62,63^. Therefore, these proteins are likely of very low abundance within platelet cells and would have been overwhelmed by more abundant proteins during LC-MS/MS analysis of platelet lysates. In contrast, the simpler releasate protein mixture exhibits a lower dynamic range which facilitates detection of minor protein components with high sensitivity. This demonstrates the advantage of platelet releasate analysis compared to analysis of lysates alone.

In conclusion, the comprehensive dataset we provide here on the platelet proteome provides the groundwork for future mechanistic studies investigating the functions of the novel proteins identified and the previously uncharacterised O-glycosylation sites. Analysis of how the platelet proteome is altered in disease states such as type II diabetes, where platelets are known to be hyperactivated^64^, should prove fruitful to detect new mechanisms of platelet dysfunction. In addition to the abundant O-glycosylation decorating the platelet releasate proteins, many other PTMs remain to be explored in the context of platelet function and hemostasis.

## ACKNOWLEDGEMENTS

M.L. is a Cancer Institute New South Wales Future Research Leader Fellow. F.H.P. is recipient of a Sydney Cardiovascular Fellowship, University of Sydney and the Heart Research Institute and a Ministry of Health NSW Cardiovascular Early Mid-Career Grant. Y.K. and P.R.C. are recipients of a Sydney Cardiovascular Initiative Catalyst Award in Precision Medicine 2020. M.T.-A. is an ARC Future Fellowship recipient (FT210100455) and supported by an MQ Enterprise Partnership Scheme. T.H.C. is supported by an International Research Training Program Scholarship from Macquarie University (20224231). This work was partially supported by a grant from the National Institute of General Medical Sciences to R.S.H. (GM061126). We thank SydneyMS for providing the instrumentation used in this study.

## AUTHORSHIP

## Contribution

C.B.H., Y.K., B.J., M.C., T.H.C., P.R.C., H.H., M.W. and M.L. performed experiments and analysed results; F.H.P., M.T.-A., R.S.H., and M.L. designed the research and wrote the paper.

