## Supplementary Material for "Ultra-sensitive platelet proteome maps the O-glycosylation landscape and charts the response to thrombin dosage"

[1]The Heart Research Institute and Charles Perkins Centre, University of Sydney, Sydney, NSW, Australia. [2]Central Clinical School, The University of Sydney, Sydney, NSW, Australia. [3]Charles Perkins Centre, School of Medical Sciences, Sydney, NSW, Australia. [4]School of Natural Sciences, Macquarie University, Macquarie Park, NSW, Australia. [5]Complex Carbohydrate Research Center, Department of Biochemistry and Molecular Biology, University of Georgia, Athens, Georgia, USA.

**Supplementary Methods**

**Patient blood collection**

Human ethics was from the University of Sydney (approval number 2014/244). Venepuncture was performed using a 19-gauge needle and light tourniquet. Blood was collected using 16 x 100 mm 8.5 mL vacutainer glass whole blood acid-citrate-dextrose (ACD) tubes (Becton Dickinson, Cat# 366645) and gently mixed following collection. Blood was collected from healthy volunteers free from medication for the past 10 days ranging from 22-60 years old (median age 32 years old).

**Platelet isolation from whole blood**

Platelets were isolated from whole blood within 3-4 hours post-venepuncture. Whole blood was fractioned by centrifugation (200 x g for 20 minutes, brake = 0) to separate platelet rich plasma (PRP) from red blood cell (RBC) and white blood cell (WBC) fractions. All centrifugation steps were performed at room temperature. PRP was rested for 30 minutes in a water bath at 37°C before the addition of 20% (v/v) pre-warmed (37°C) acid citrate-dextrose (ACD) (Cat# C3821, Sigma-Aldrich). Platelets were separated from plasma by centrifugation (800 x g for 10 minutes, brake = 4). The platelets were resuspended in pre-warmed (37°C) modified HEPES/Tyrodes (HTGlc) buffer (129 mM NaCl, 0.34 mM $Na_2HPO_4$, 2.9 mM KCl, 12 mM $NaHCO_3$, 20 mM HEPES, 5 mM glucose, 1 mM $MgCl_2$; pH 7.4). Resuspended platelets were rested for 20 minutes in a 37°C water bath following the addition of 10% (v/v) pre-warmed (37°C) ACD and 0.02 U/mL apyrase. Platelets were pelleted by centrifugation (800 x g for 5 minutes, brake = 4) before resuspension in prewarmed HTGlc buffer at the working concentration of 400 x $10^3$/µL. Addition of prostaglandin E1 (PGE1) (2 µM) took place immediately before all centrifugation steps to minimize platelet activation. Platelet concentration was maintained below $10^6$ / µL during washing stages. Platelet concentration was measured using a Sysmex KX-21N haemocytometer.

**Separation of platelet releasate and lysate for proteomics**

Washed platelets (400 x $10^3/\mu L$) were divided into 250 µL aliquots. To a resting control, 0.02 U/mL apyrase was added, while other aliquots were activated with thrombin (Sigma, Cat# T4648) at a final activity of 0.2 U/mL for maximal stimulation or 0.025 U/mL for submaximal stimulation. All samples were incubated in a 37°C water bath for 5 min. Following incubation, PPACK (25 nM) was added to the thrombin-stimulated sample and 2 µM PGE1 was added immediately prior to centrifugation of the resting sample. The supernatant at this point was regarded as the "platelet releasate" and was aspirated and stored under argon at -80°C. The remaining platelet protein was regarded as the "platelet lysate". Pellet lysate was obtained via resuspension in SDC lysis buffer (4% (w/v) sodium deoxycholate in 0.1 M Tris-HCl pH 8.0) and heating at 95°C for 10 min. Protein concentration was determined by BCA assay (Thermo Fisher). Platelet lysates were aliquoted and stored under argon at -80°C.

**Flow cytometry analysis of washed platelet contamination and platelet pre-activation**
The expression of key platelet membrane-specific proteins $\alpha_{IIb}\beta_3$ and P-selectin (CD62p) was used to assess platelet pre-activation prior to thrombin stimulation. Platelet pre-activation was measured following platelet releasate and lysate storage and within 5 hours post-venepuncture. Washed platelets were suspended in HTGlc buffer at a concentration of 10 x $10^3/\mu L$ and stained with either mouse anti-human $\alpha_{IIb}\beta_3$ IgG conjugated with FITC (Becton Dickinson Biosciences, Cat# 340507) or mouse anti-human CD62p IgG conjugated with APC (Cat# 550888). Platelet activation was achieved via the addition of thrombin (0.025 or 0.2 U/mL) to washed platelet preparation for 10 minutes at room temperature and was cancelled via the addition of 25 µM D-phenylalanyl-N-[(1S)-4-[(aminoiminomethyl)amino]-1-(2-chloroacetyl) butyl]-L-prolinamide dihydrochloride (PPACK). Washed unstained, stained resting, and stained thrombin-activated platelets were analysed by flow cytometry using a Becton Dickinson Accuri C6 flow cytometer to confirm expression of $\alpha_{IIb}\beta_3$ and CD62p. Data analysis was achieved using FlowJo software (FlowJo, LLC).

For analysis of WBC and RBC contamination in washed platelet preparations, samples were stained with anti-human CD45 IgG conjugated to PerCP-Cy™5.5 (Becton Dickinson Biosciences, Cat# 340953,) and mouse anti-human RBC IgG conjugated to FITC (Cat# 349103) respectively. Both samples were additionally stained with mouse anti-human CD41 IgG conjugated with FITC (Cat# IMO649U). Unstained washed platelets were used as the control and cell fluorescence was recorded by flow cytometry using a Accuri C6 flow cytometer. CD41+ cells were gated as platelets (**Figure 3a**). Since platelets are significantly smaller in size than RBCs and WBCs and have significantly less granularity than WBCs, events beyond the upper limits of platelet side scatter (granularity) and forward scatter (size) were regarded as either WBCs or RBCs (**Figure 3a**). Events above these upper limits which were CD45+ were regarded as WBCs, and RBC+ events were regarded as RBCs. Event counts of each sub-population were used to derive a ratio of platelets to WBCs and platelets to RBCs at a higher sensitivity than which was achievable using a cell counter.

**Determination of flow cytometry cut-offs for pre-activation and thrombin doses**
The laboratory has an established routine platelet isolation protocol by sequential centrifugation steps as described previously[1] and detailed below. Using this method, we have determined the % staining for mouse anti-human CD62p IgG conjugated with APC (Cat# 550888) and anti-human αIIbβ3 IgG (PAC-1) conjugated with FITC (Becton Dickinson

Biosciences, Cat# 340507) in the resting platelet population compared with unstained resting platelets (n=30). PAC-1 detects the active confirmation of αIIbβ3. The average % + 2 SD of this historical group was used as a cut-off for inclusion of resting platelet preparations from healthy donors included in this study. Samples that exhibited CD62P >25% and PAC-1 >15% were excluded from further analysis. These percentages for CD62P and PAC-1 are in line with a previous study of platelet isolation for proteomic analysis[2].

For establishing sub-maximal ("low") and maximal ("high") thrombin dose, platelets were isolated from 5 healthy donors. Platelet aggregation was recorded over 10 min to doses of thrombin: 0.02, 0.03, 0.05 and 0.2 U/ml. LTA was evaluated as described using an AggRAM 1484[3]. Experiments were conducted in 300 µL aliquots of washed platelets (400 x $10^3$/µL) buffered in HTGlc. The washed platelet suspension was mixed by magnetic stirrer. After the addition of thrombin, platelet aggregation was recorded for 10 minutes. The maximum aggregation was determined as the peak light transmission. Based on the aggregation response for this group of individuals and the batch of thrombin used in this study, we determined submaximal (low) dose of thrombin as 0.025 U/ml and maximal (high) dose of thrombin as 0.2 U/ml.

**SDS-PAGE and Coomassie staining**

Protein samples were reduced and denatured in SDS and beta-mercaptoethanol (b-ME) at 95°C for 10 minutes before loading onto pre-cast 4–20% polyacrylamide gels (Cat# 4561094, Bio Rad Laboratories). Electrophoresis was performed for 1.5 h at a voltage of 100 V alongside Novex pre-stained protein standards (Cat# LC5800, Invitrogen). Following electrophoresis, gels stained with Coomassie Brilliant Blue R-250 Staining Solution (Cat# 161-0436, Bio Rad Laboratories) for 1 hour with gentle agitation. Gels were imaged using a near infrared fluorescence scanner (Odyssey CLx Imaging System, Li-Cor).

**O-Glycome sample preparation**

The O-glycans were chemically released from the mixture of proteins extracted from platelet releasates after complete enzymatic elimination of any attached *N*-glycans, and the liberated *O*-glycans were processed as previously described[4]. Bovine fetuin (Sigma-Aldrich) was included as a sample handling and LC-MS/MS control. Briefly, 20 µg total protein from each platelet releasate sample (and from bovine fetuin) was reduced with 10 mM aqueous dithiothreitol (DTT) for 45 min at 56°C and carbamidomethylated with 25 mM aqueous iodoacetamide for 30 min in the dark at 20°C. The alkylation reaction was quenched with 30 mM aqueous DTT (final concentrations stated). The proteins were spotted onto an activated 0.45 µm PVDF membrane (Merck-Millipore), dried, stained with Direct Blue, and excised. The excised spots were transferred to separate wells in a flat bottom polypropylene 96-well plate (Corning Life Sciences, Melbourne, Australia), blocked with 1% (w/v) polyvinylpyrrolidone in 50% (v/v) aqueous methanol, and washed with MilliQ water. The *N*-glycans were exhaustively released using 2 U recombinant *Elizabethkingia miricola* peptide:*N*-glycosidase F (PNGase F) expressed in *Escherichia coli* (Promega) per 20 µg protein in 10 µL water per well and incubated for 16 h at 37°C. A second round of PNGase F-based *N*-glycan release was performed the next day to ensure complete removal of all *N*-glycans from the protein samples to avoid cross-contamination of *N*-glycans in the subsequent *O*-glycan samples. The *O*-glycans were subsequently released by incubation with 20 µL 0.5 M sodium borohydride in 50 mM aqueous potassium hydroxide for 16 h at 50°C. The reduction reaction was then quenched using 2 µL glacial acetic acid and the released and reduced *O*-glycans were transferred into fresh 1.5 mL Eppendorf tubes. Dual desalting of the reduced *O*-glycans was performed using firstly strong cation exchange resin (AG 50W-X8 Resin, Bio-Rad) (where the *O*-glycans were not retained), followed by porous graphitised carbon (PGC) resin (where *O*-glycans were retained) custom packed as micro-columns on top of C18 discs (Merck-Millipore) in P10 solid-phase extraction (SPE) formats. Following micro-column equilibration and sample loading and washing, the *O*-glycans were eluted from the PGC-SPE micro-columns using 0.05% trifluoroacetic acid/40% acetonitrile (ACN)/59.95% water (all v/v), dried and resuspended in 20 μL water. Samples were centrifuged at 14,000 × *g* for 10 min at 4ºC and the clear supernatant fractions were carefully transferred to high recovery glass vials (Waters) to avoid debris and particulates in the LC-MS/MS injection vials.

**O-Glycan profiling with PGC-LC-MS/MS**

The *O*-glycans were profiled using a well-established PGC-LC-MS/MS method[4,5]. In brief, the *O*-glycan samples were injected on a HyperCarb KAPPA PGC-LC column (particle/pore size, 3 μm/250 Å; column length, 30 mm; inner diameter, 0.181 mm, Thermo Hypersil, Runcorn, UK) heated to 50ºC. The *O*-glycans were separated over a 60 min linear gradient of 0–45% (v/v) pure ACN (solvent B) in 10 mM aqueous ammonium bicarbonate (solvent A) on a 1260 Infinity Capillary HPLC system (Agilent) operating with a constant flow rate of 20 μL/min. The separated *O*-glycans were introduced directly into the mass spectrometer, ionised using electrospray ionisation and detected in negative ion polarity mode using a linear trap quadrupole Velos Pro ion trap mass spectrometer (Thermo Fisher Scientific). The acquisition settings included a full MS1 scan acquisition range of *m/z* 300–2000, resolution of *m/z* 0.25 full width half maximum and a source voltage of +3.2 kV. The automatic gain control for the MS1 scans was set to $5 \times 10^4$ with a maximum accumulation time of 50 ms. For the MS/MS events, the resolution was set to *m/z* 0.25 full width half maximum, the automatic gain control was $2 \times 10^4$ and the maximum accumulation time was 300 ms. Data-dependent acquisition was enabled for all samples. The three most abundant precursors in each MS1 full scan were selected for fragmentation using resonance activation (ion trap) collision-induced dissociation (CID) at a normalised collision energy of 33%. Dynamic exclusion of precursors was inactivated. All MS and MS/MS data were acquired in profile mode. The mass accuracy of the precursor and product ions was typically better than 0.2 Da. The LC-MS/MS instrument was tuned and calibrated, and its performance bench marked using well-characterised bovine fetuin *O*-glycan standards analysed at the same time as the samples of interest. The generated LC-MS/MS raw data files (made publicly available via GlycoPOST[6], accession number GPST000211) were browsed, interrogated, and manually annotated using Xcalibur v2.2 (Thermo Fisher Scientific), GlycoMod[7] and GlycoWorkBench v2.1[8] as previously described[9]. Briefly, glycans were identified based on the monoisotopic precursor mass, the match between the observed and theoretical MS/MS fragmentation pattern *in silico* generated using GlycoWorkBench, and the relative and absolute PGC-LC retention time of each glycan. Additional support for some structures was obtained using PGC-LC retention time matching of observed platelet *O*-glycans to known bovine fetuin *O*-glycans[10]. Further, the reported platelet *O*-glycan structures were backed by observations of identical or similar *O*-glycan structures in the mammalian glycobiology literature. The relative abundances of the confidently identified *O*-glycans were determined from area-under-the-curve (AUC) measurements based on extracted ion chromatograms performed for all relevant charge states of the monoisotopic precursor *m/z* using Xcalibur v2.2 (Thermo Fisher Scientific).

**Production of MMRN1 in HEK293T WT, *POFUT1* KO and *POFUT2* KO cells**

HEK293T WT, *POFUT1* KO[11], or *POFUT2* KO[12] cells were transiently transfected with pcDNA3.1-hMMRN1-Myc-His6 (10 μg/plate) in 10 x 10-cm dishes containing 8 ml/plate

Opti-MEM (Invitrogen) using 60 µl PEI/plate. Two days later, the media from the 10 plates of cells was combined. MMRN1 protein was purified using Ni-NTA agarose (Qiagen) and eluted with 600 µl of 250 mM imidazole as described previously[13]. 10 µl of the input media, flowthrough, first elution and second elution were collected and loaded onto 4-20% SDS-PAGE (Invitrogen), transferred to a nitrocellulose membrane. The membrane was incubated with anti-Myc antibody (Clone 9E10, Invitrogen, 1:2500) and anti-His antibody (Clone AD1.1.10, BIO-RAD, 1:1500), and subsequently with IDRye 800-conjugated goat anti-mouse IgG antibody (LI-COR, 1:2500). The Western blot bands were visualized using Odyssey System (LI-COR).

**Mass spectrometry of MMRN1 produced in HEK293T WT, *POFUT1* KO and *POFUT2* KO cells**

Purified MMRN1 (400 µl out of 600 µl Ni-NTA elution) from HEK293T WT, *POFUT1* KO or *POFUT2* KO cells was acetone precipitated, reduced, alkylated and digested with trypsin as described[13]. Peptides were analysed by LC-MS/MS using an EasyNano-LC with a C18 EasySpray PepMap RSLC C18 column (50 µm X 15 cm, Thermo) coupled with a Thermo Fisher Q-Exactive Plus mass spectrometer. Data analysis was performed with Byonic (Protein Metrics). A suggested search by Preview (Protein Metrics) was performed to set up the basic search parameters including cleavage specificity, mass tolerance, fixed and variable modifications. Glycans were set as variable modifications (common1) using a customed O-glycan search space including Fuc(1), Hex(1)Fuc(1), HexNAc(1)Fuc(1), Hex(1) and HexNAc(1). To make the extracted ion chromatograms (EICs) for a given peptide, the ions of each glycoform were extracted from the MS1 spectrum using their respective m/z, then overlaid to compare the relative ion intensity. The EICs were smoothed using Gauss algorithm.

**Supplementary Tables and Files**

**Supplementary Table 1**. Platelet lysate and releasate proteome quantification data table.

**Supplementary Table 2**. Peptide identification data for the open search detection of modified platelet releasate proteins.

**Supplementary Table 3**. O-Glycome profiling of platelet releasate proteins.

**Supplementary Table 4**. Peptide identification data for EThcD analysis of platelet releasate proteins.

**Supplementary Table 5**. Platelet releasate proteins significantly increased after thrombin stimulation.

**Supplementary File 1**. Platelet releasate database for open search with decoys.

**Supplementary File 2**. PGC-LC-MS/MS-based O-Glycan characterisation including annotated spectra and peak quantification.

**Supplementary Methods References**


1. Cazenave JP, Ohlmann P, Cassel D, Eckly A, Hechler B, Gachet C. Preparation of washed platelet suspensions from human and rodent blood. *Methods Mol Biol*. 2004;272:13-28.
2. Wrzyszcz A, Urbaniak J, Sapa A, Wozniak M. An efficient method for isolation of representative and contamination-free population of blood platelets for proteomic studies. *Platelets*. 2017;28(1):43-53.
3. Hayward CP, Moffat KA, Pai M, et al. An evaluation of methods for determining reference intervals for light transmission platelet aggregation tests on samples with normal or reduced platelet counts. *Thromb Haemost*. 2008;100(1):134-145.
4. Jensen PH, Karlsson NG, Kolarich D, Packer NH. Structural analysis of N- and O-glycans released from glycoproteins. *Nat Protoc*. 2012;7(7):1299-1310.
5. Hinneburg H, Chatterjee S, Schirmeister F, et al. Post-Column Make-Up Flow (PCMF) Enhances the Performance of Capillary-Flow PGC-LC-MS/MS-Based Glycomics. *Anal Chem*. 2019;91(7):4559-4567.
6. Watanabe Y, Aoki-Kinoshita KF, Ishihama Y, Okuda S. GlycoPOST realizes FAIR principles for glycomics mass spectrometry data. *Nucleic Acids Research*. 2020;49(D1):D1523-D1528.
7. Cooper CA, Gasteiger E, Packer NH. GlycoMod--a software tool for determining glycosylation compositions from mass spectrometric data. *Proteomics*. 2001;1(2):340-349.
8. Ceroni A, Maass K, Geyer H, Geyer R, Dell A, Haslam SM. GlycoWorkbench: a tool for the computer-assisted annotation of mass spectra of glycans. *J Proteome Res*. 2008;7(4):1650-1659.
9. Hinneburg H, Pedersen JL, Bokil NJ, et al. High-resolution longitudinal N- and O-glycoprofiling of human monocyte-to-macrophage transition. *Glycobiology*. 2020;30(9):679-694.
10. Kozak RP, Urbanowicz PA, Punyadeera C, et al. Variation of Human Salivary O-Glycome. *PLoS One*. 2016;11(9):e0162824.
11. Takeuchi H, Yu H, Hao H, et al. O-Glycosylation modulates the stability of epidermal growth factor-like repeats and thereby regulates Notch trafficking. *J Biol Chem*. 2017;292(38):15964-15973.
12. Benz BA, Nandadasa S, Takeuchi M, et al. Genetic and biochemical evidence that gastrulation defects in Pofut2 mutants result from defects in ADAMTS9 secretion. *Dev Biol*. 2016;416(1):111-122.
13. Kakuda S, Haltiwanger RS. Analyzing the posttranslational modification status of Notch using mass spectrometry. *Methods Mol Biol*. 2014;1187:209-221.

### Supplementary Figure 1

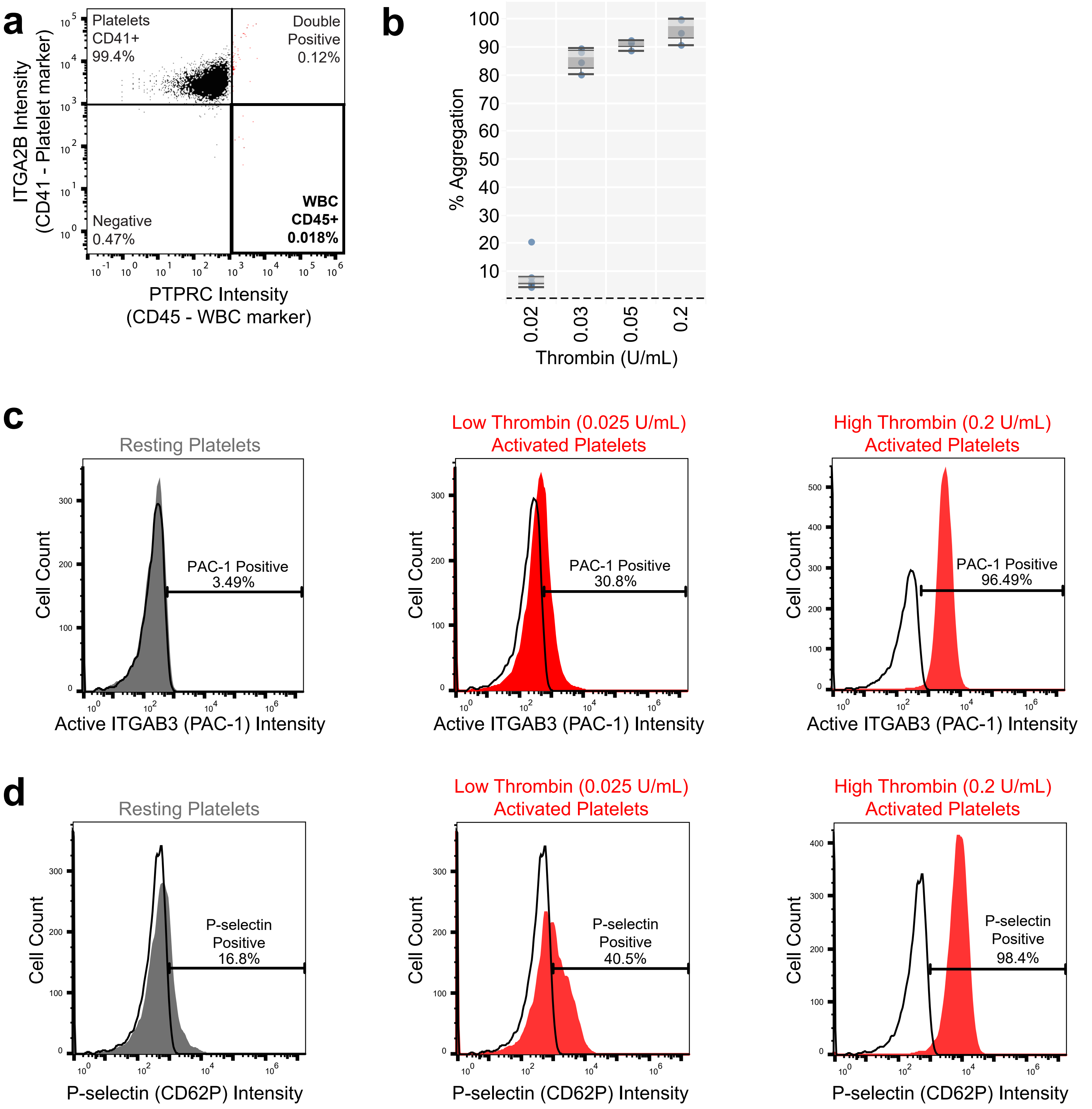


**Supplementary Figure 1. Isolation of platelets in either the resting or thrombin-activated state for generation of platelet lysates and releasates.** (a) Scatterplot of flow cytometry analysis for platelet contamination by white blood cells (WBC). (b) Analysis of platelet activation with a dose response of thrombin using aggregometry analysis. (c) Histograms of platelet activation using PAC-1 intensity (x-axis). Resting platelets are shown in grey, platelets stimulated with either 0.025 or 0.2 U/mL thrombin shown in red. (d) Histograms of platelet activation using P-selectin intensity (x-axis). Resting platelets are shown in grey, platelets stimulated with either 0.025 or 0.2 U/mL thrombin shown in red.

### Supplementary Figure 2

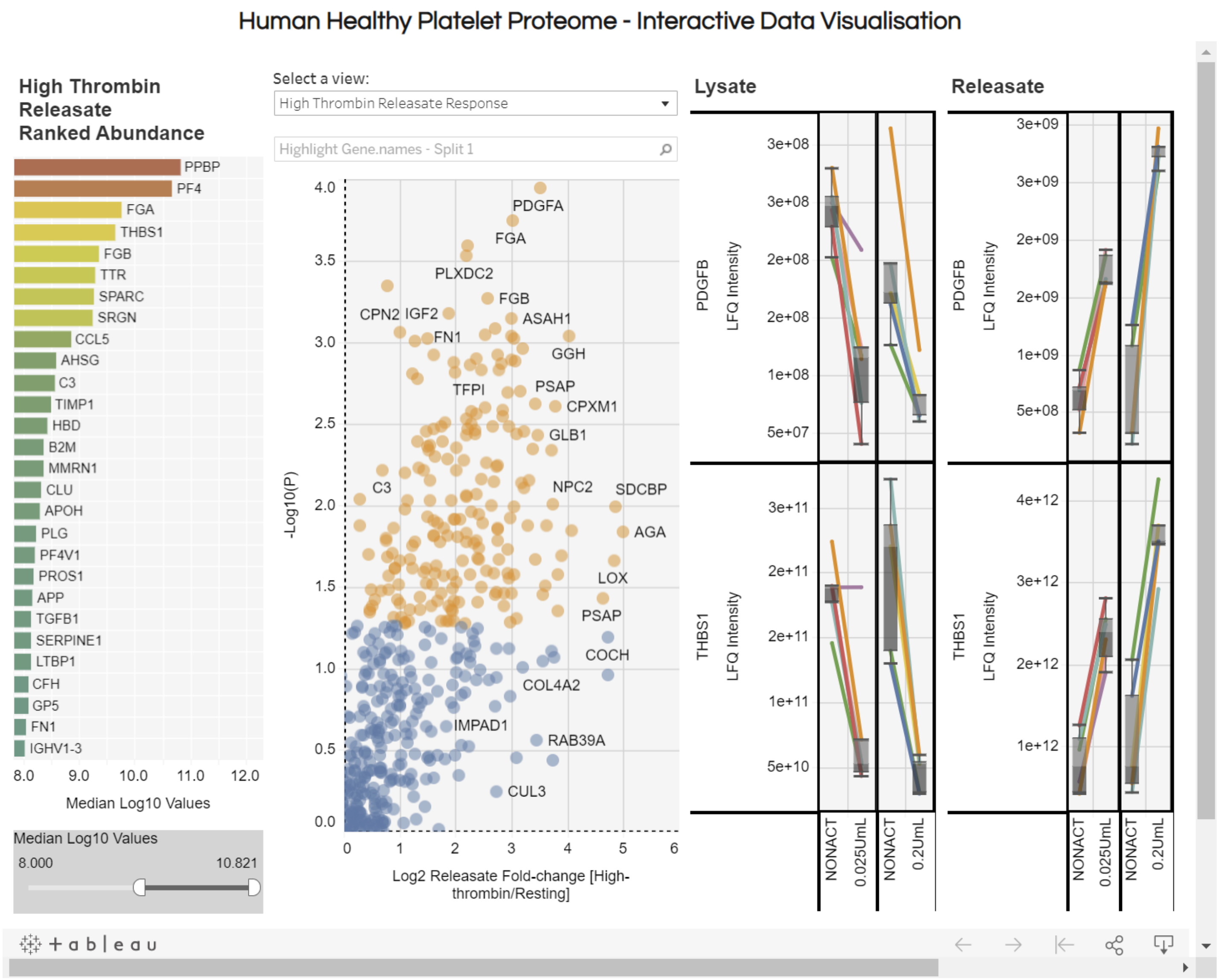


**Supplementary Figure 2. Screenshot of the platelet proteome interactive visualisation.** Using this online resource available at (larancelab.com/platelet-proteome) users can browse and analyse human platelet proteins and examine the quantitiative data generated for the response to thrombin dose.
