## Supplementary File 2 for "Ultra-sensitive platelet proteome maps the O-glycosylation landscape and charts the response to thrombin dosage"

EICs and manually annotated PGC-LC-ESI-CID-MS (-)
spectra of *O*-glycans of platelet releasate activated with thrombin (0.2 U/mL)

| Glycan # | Generic glycan composition [likely glycan structure if support available] | Glycan structure without linkage annotation | Glycan structure with linkage annotation if support available | Main experimental evidence for drawn structure | Literature further supporting depicted glycan structure |
|---|---|---|---|---|---|
| **1** | (Hex)1(Deoxyhexose)1<br>[Glc(β1-3)-Fuc] | 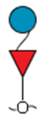 | 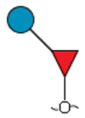 | MS,<br>MS/MS | https://pubmed.ncbi.nlm.nih.gov/30690220/ |
| **2** | (Hex)1(Pent)2<br>[Xyl(α1-3)Xyl(α1-3)Glc] | 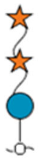 | 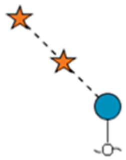 | MS,<br>MS/MS | https://academic.oup.com/glycob/article/20/3/287/1987743<br>https://www.nature.com/articles/s41419-020-03314-y.pdf<br>https://doi.org/10.1038/nrm3383 |
| **3a** | (Hex)1(HexNAc)1(NeuAc)1<br>[NeuAc(α2-3)Gal(β1-3)GalNAc] | 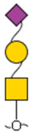 | 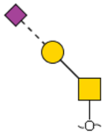 | RT matching*,<br>MS,<br>MS/MS | https://www.ludger.com/docs/products/clib/cofas/clibo-fetuin-01-b4be-03-cofa.pdf<br>https://doi.org/10.1371/journal.pone.0162824<br>https://doi.org/10.1016/j.jprot.2014.05.022 |
| **3b** | (Hex)1(HexNAc)1(NeuAc)1<br>[NeuAc(α2-3)[Gal(β1-3)]GalNAc] | 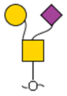 | 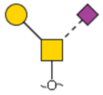 | RT matching*,<br>MS<br>MS/MS | |
| **4** | (Hex)1(HexNAc)2(NeuAc)1<br>[NeuAc(α2-3)Gal(β1-3)[GlcNAc(β1-6)]GalNAc] | 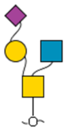 | 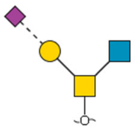 | MS,<br>MS/MS | https://doi.org/10.1016/j.chom.2020.08.004 |
| **5** | (Hex)1(HexNAc)1(NeuAc)2<br>[NeuAc(α2-3)Gal(β1-3)(NeuAcα2-6)GalNAc] | 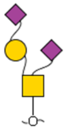 | 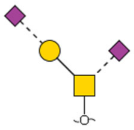 | RT matching*,<br>MS,<br>MS/MS | https://www.ludger.com/docs/products/clib/cofas/clibo-fetuin-01-b4be-03-cofa.pdf<br>https://doi.org/10.1371/journal.pone.0162824<br>https://doi.org/10.1016/j.jprot.2014.05.022 |

| | | | | | |
|---|---|---|---|---|---|
| **6a** | (Hex)2(HexNAc)2(NeuAc)1<br>[Gal(β1-4)GlcNAc(β1-6)[NeuAc(α2-3)Gal(β1-3)]GalNAc] | 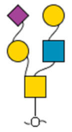 | 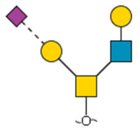 | RT matching**,<br>MS,<br>MS/MS | https://doi.org/10.1016/S0021-9258(18)41768-1<br>https://doi.org/10.1016/S0021-9258(17)32238-X<br>https://doi.org/10.1074/jbc.M202921200<br>https://doi.org/10.1093/glycob/cwaa020 |
| **6b** | (Hex)2(HexNAc)2(NeuAc)1<br>[NeuAc(α2-3)Gal(β1-4)GlcNAc(β1-6)[Gal(β1-3)]GalNAc] | 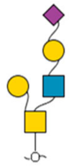 | 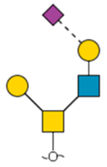 | RT matching**,<br>MS<br>MS/MS | |
| **7** | (Hex)2(HexNAc)2(Deoxyhexose)1(NeuAc)1<br>[NeuAc(α2-3)Gal(β1-4)[Fuc(α1-3)]GlcNAc(β1-6)[Gal(β1-3)]GalNAc] | 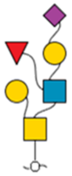 | 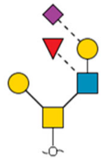 | MS,<br>MS/MS | https://doi.org/10.1021/bi972612a<br>https://doi.org/10.1093/glycob/4.2.203 |
| **8** | (Hex)1(HexNAc)1(NeuAc)3 | 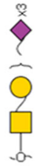 | 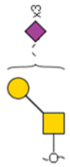 | MS,<br>MS/MS | https://pubs.acs.org/doi/10.1021/acs.analchem.0c01794 |
| **9** | (Hex)2(HexNAc)2(NeuAc)2<br>[NeuAc(α2-3)Galβ1-3(NeuAc(α2-3)Gal(β1-4)GlcNAc(β1-6))GalNAc] | 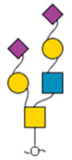 | 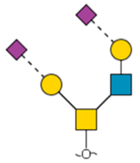 | RT matching*,<br>MS,<br>MS/MS | https://www.ludger.com/docs/products/clib/cofas/clibo-fetuin-01-b4be-03-cofa.pdf<br>https://doi.org/10.1371/journal.pone.0162824<br>https://doi.org/10.1016/j.jprot.2014.05.022 |

* Retention time matching was performed against O-glycan samples of the well characterised bovine fetuin analysed on the same PGC-LC gradient acquired in the same batch as the platelet samples of interest.[1]
** Retention time comparison and spectral matching were performed against O-glycan samples of human monocytes and macrophages analysed on a similar PGC-LC-MS/MS setup but in a different period.[2]

### Symbol, linkage and fragmentation key

- Fucose (Fuc)
- Galactose (Gal)
- Glucose (Glc)
- *N*-Acetylglucosamine (GlcNAc)
- *N*-Acetylgalactosamine (GalNAc)
- *N*-Acetylneuraminic acid (NeuAc)
- Xylose (Xyl)

Unknown linkage

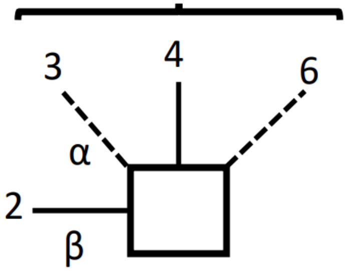


- Indicates mostly Y ions (includes oxygen of glycosidic linkage)
- Indicates mostly Z ions (excludes oxygen of glycosidic linkage)
- Reduced reducing end
- a-b Isomers

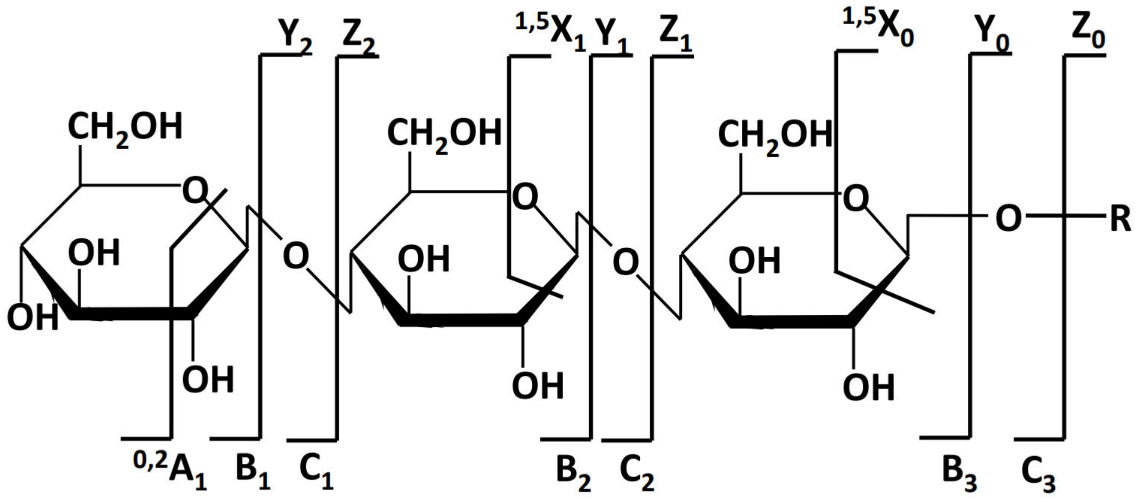


Han, L. & Costello, C.E. Mass spectrometry of glycans. *Biochemistry (Moscow)* 78, 710-720 (2013)

**Glycan #1**
**Extracted ion chromatogram**
(*m/z* 327.12)

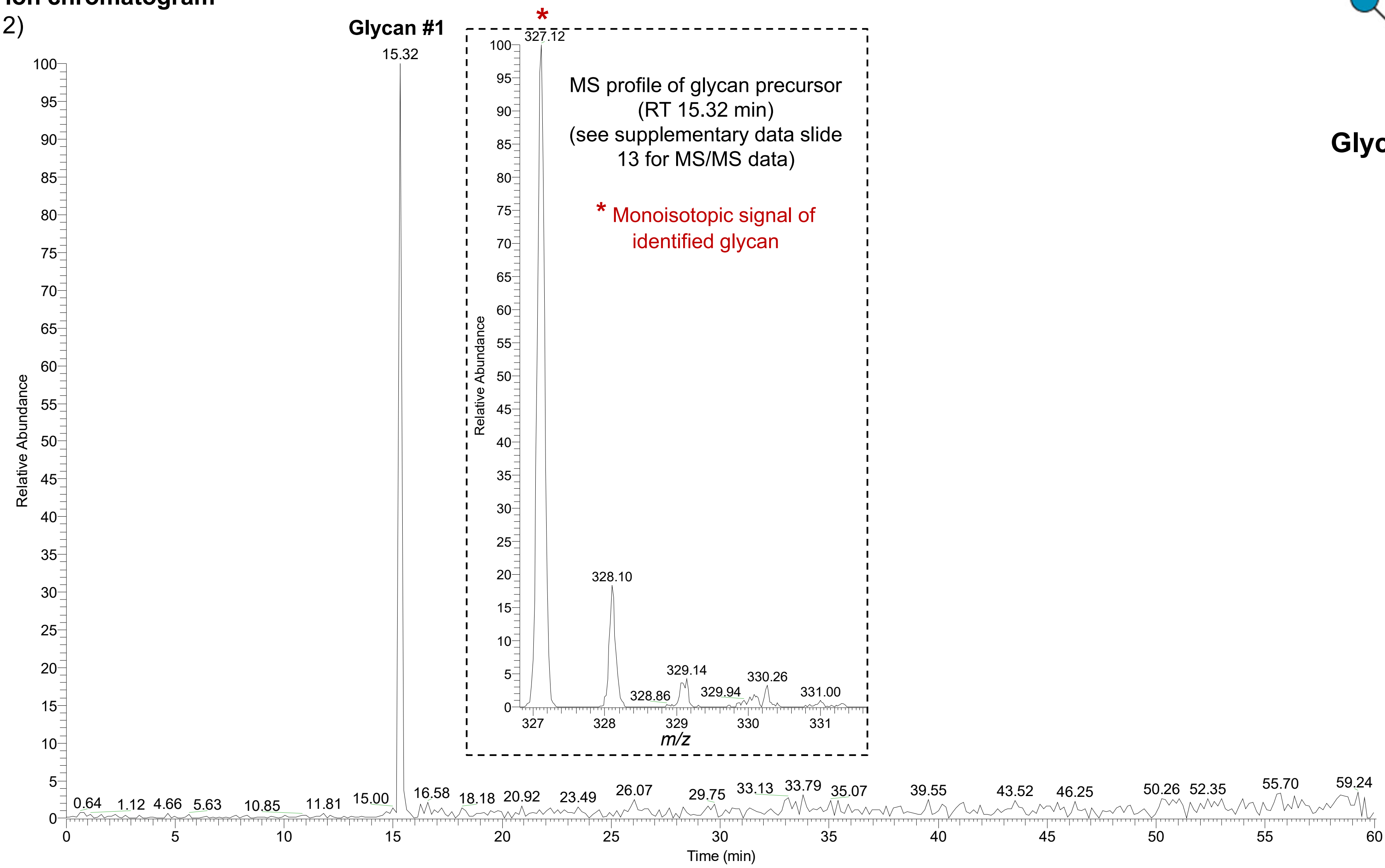


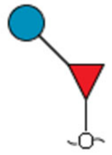

**Glycan #1**

**Glycan #2**
**Extracted ion chromatogram**
(*m/z* 445.10)

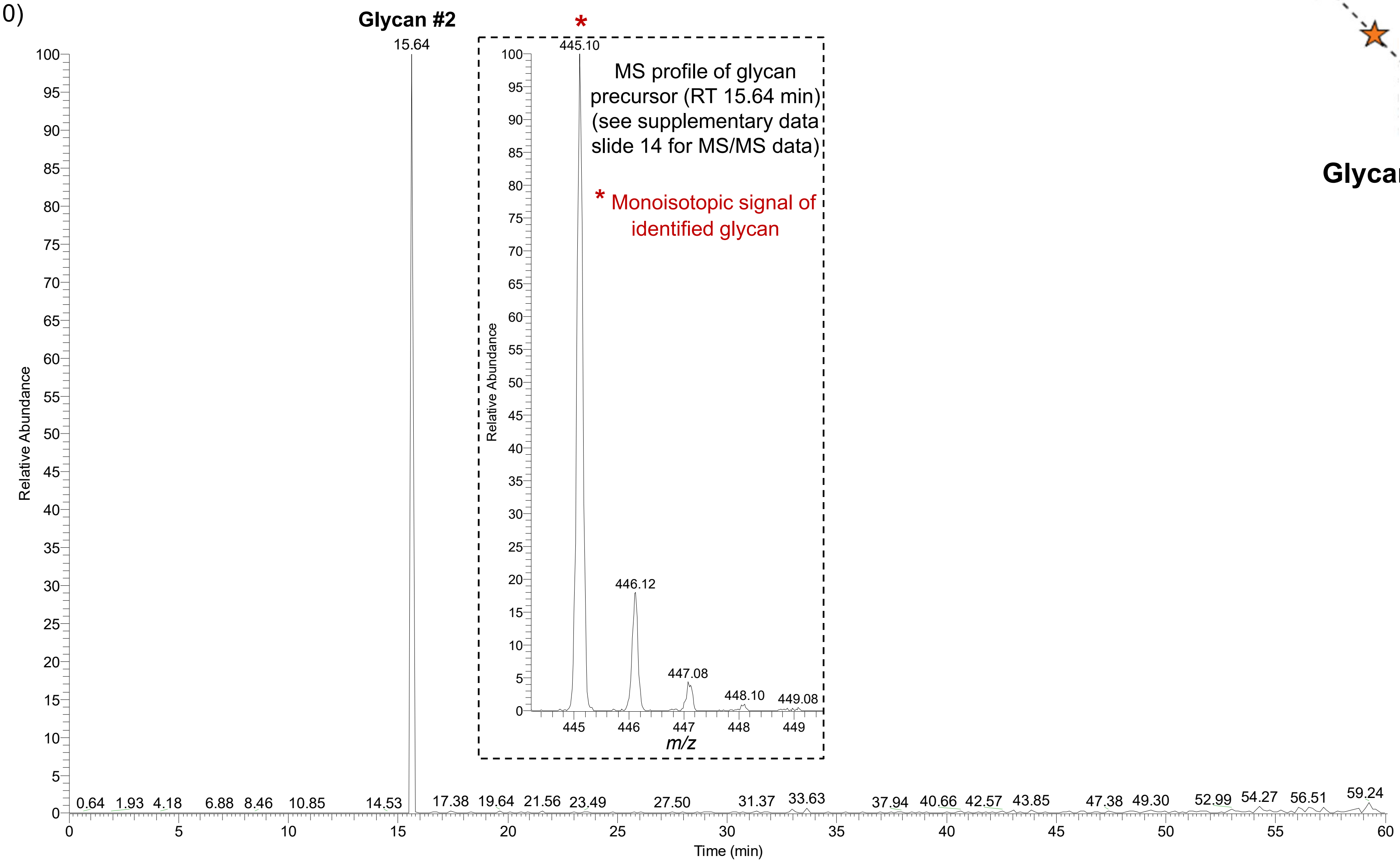


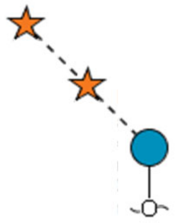

**Glycan #2**

**Glycan #3**
**Extracted ion chromatogram**
(*m/z* 675.16)

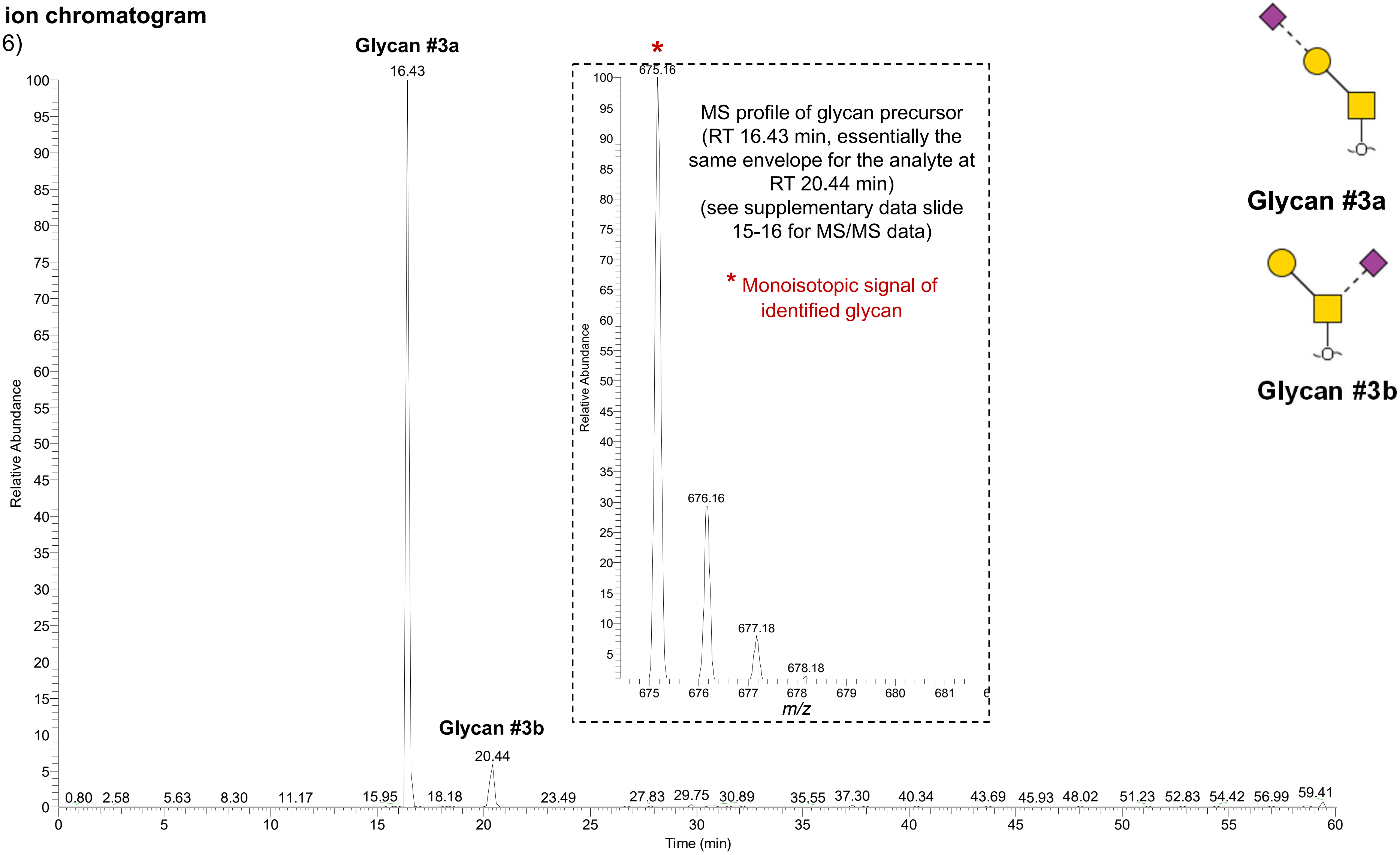

**Glycan #4**
**Extracted ion chromatogram**
($m/z$ 878.22)

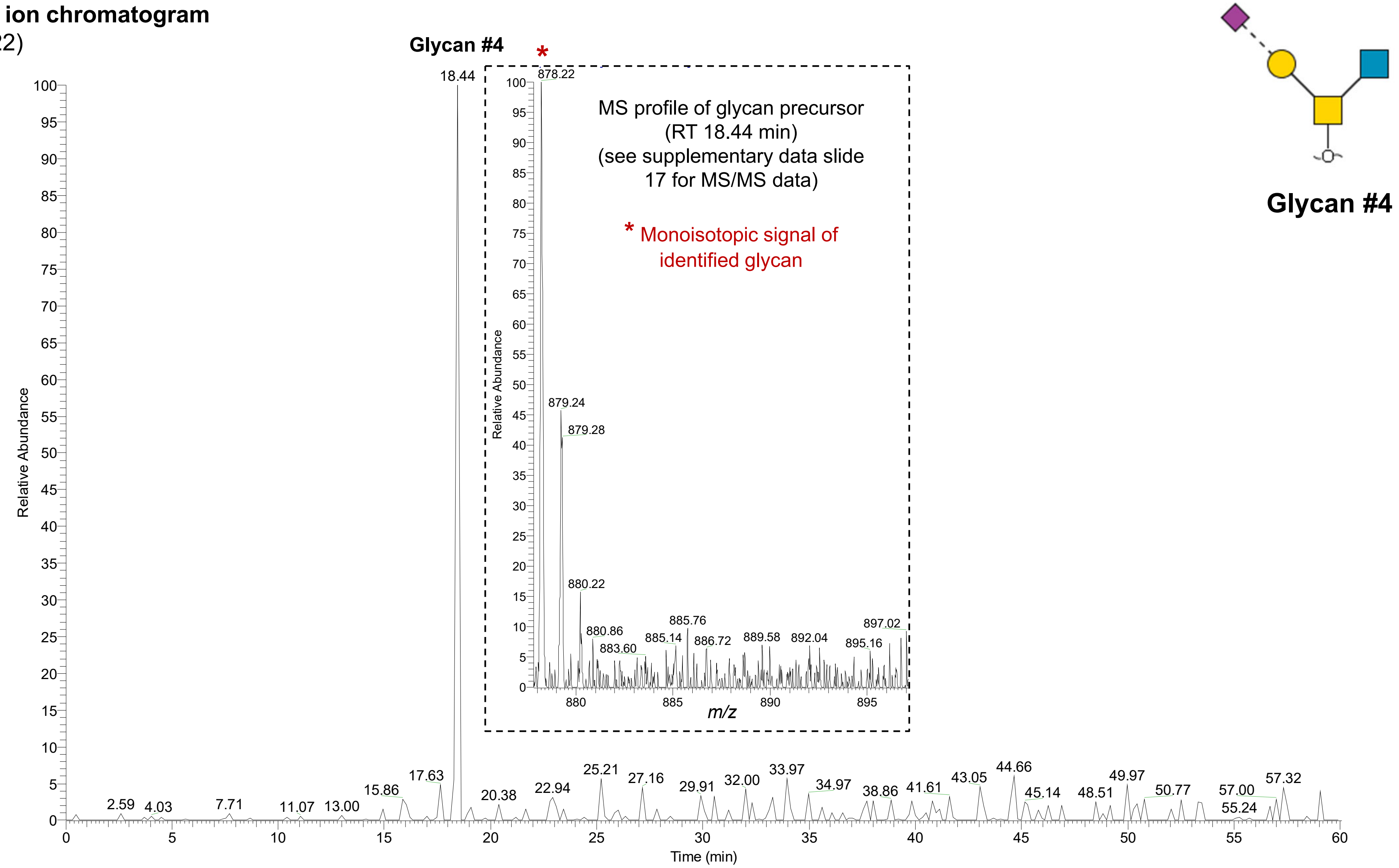

**Glycan #5**
**Extracted ion chromatogram**
(*m/z* 966.26)

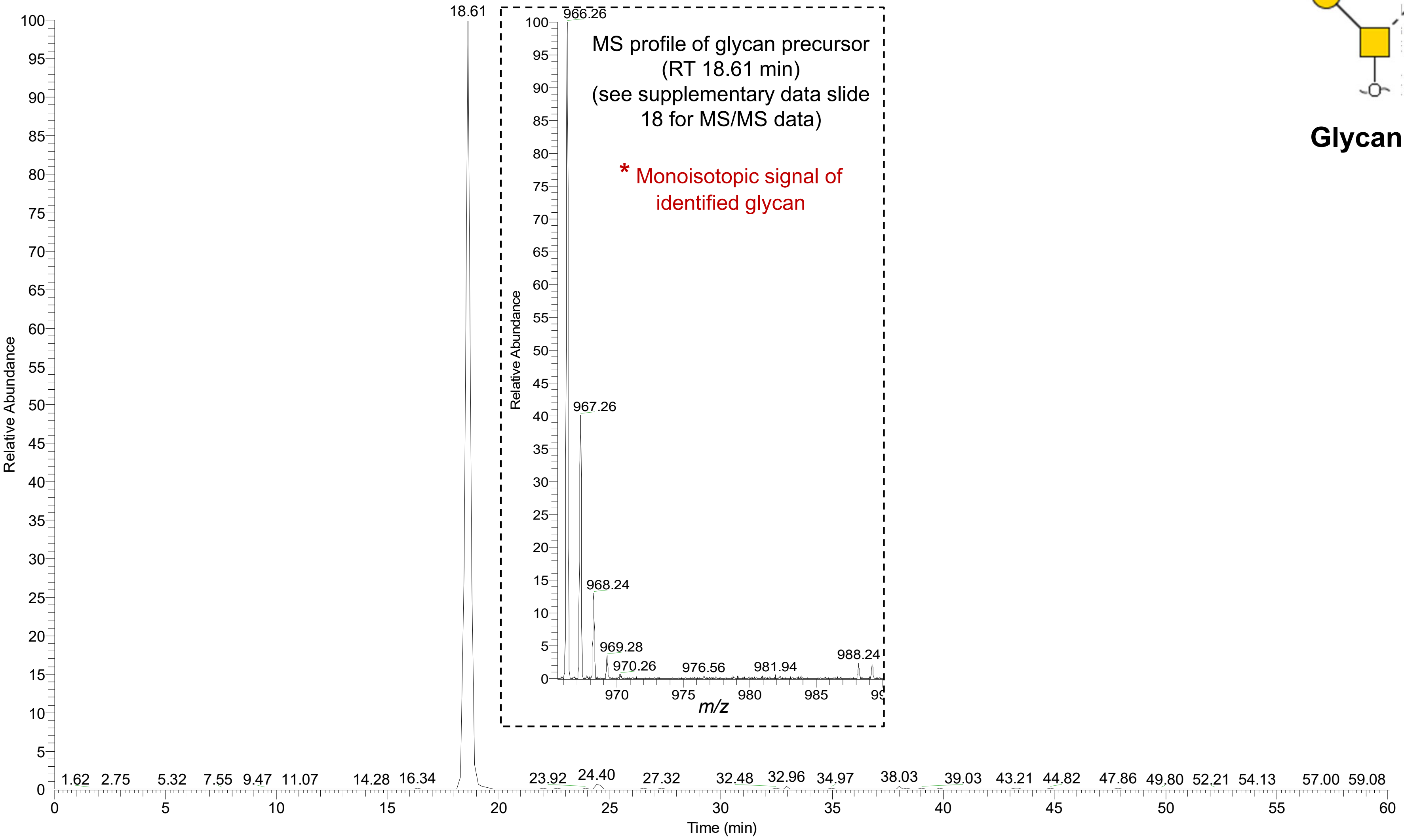


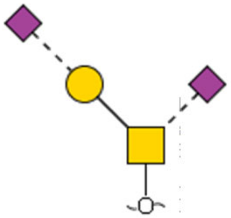

**Glycan #5**

**Glycan #6**
**Extracted ion chromatogram**
(*m/z* 1040.32)

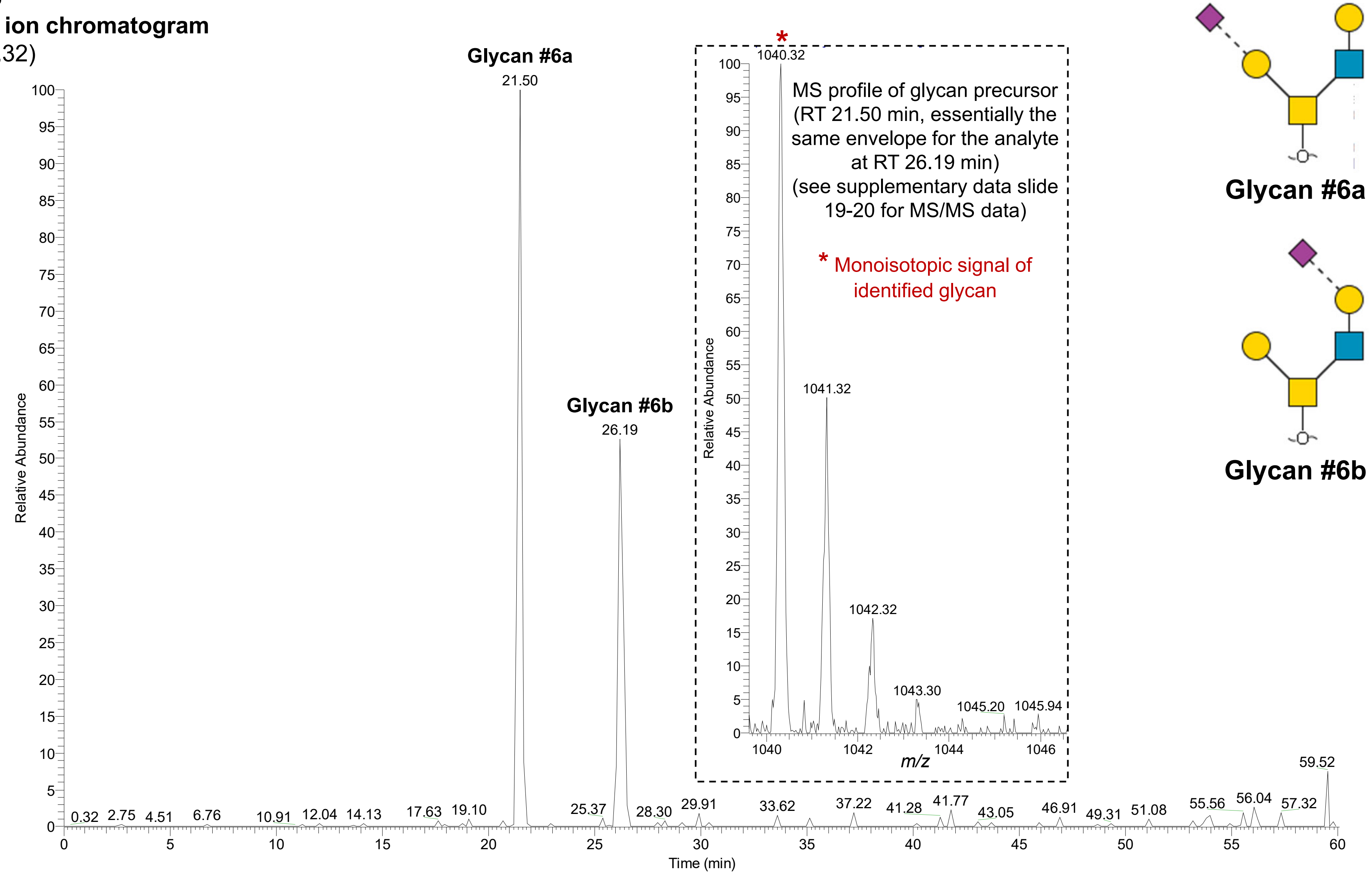

**Glycan #7**
**Extracted ion chromatogram**
(*m/z* 1186.38)

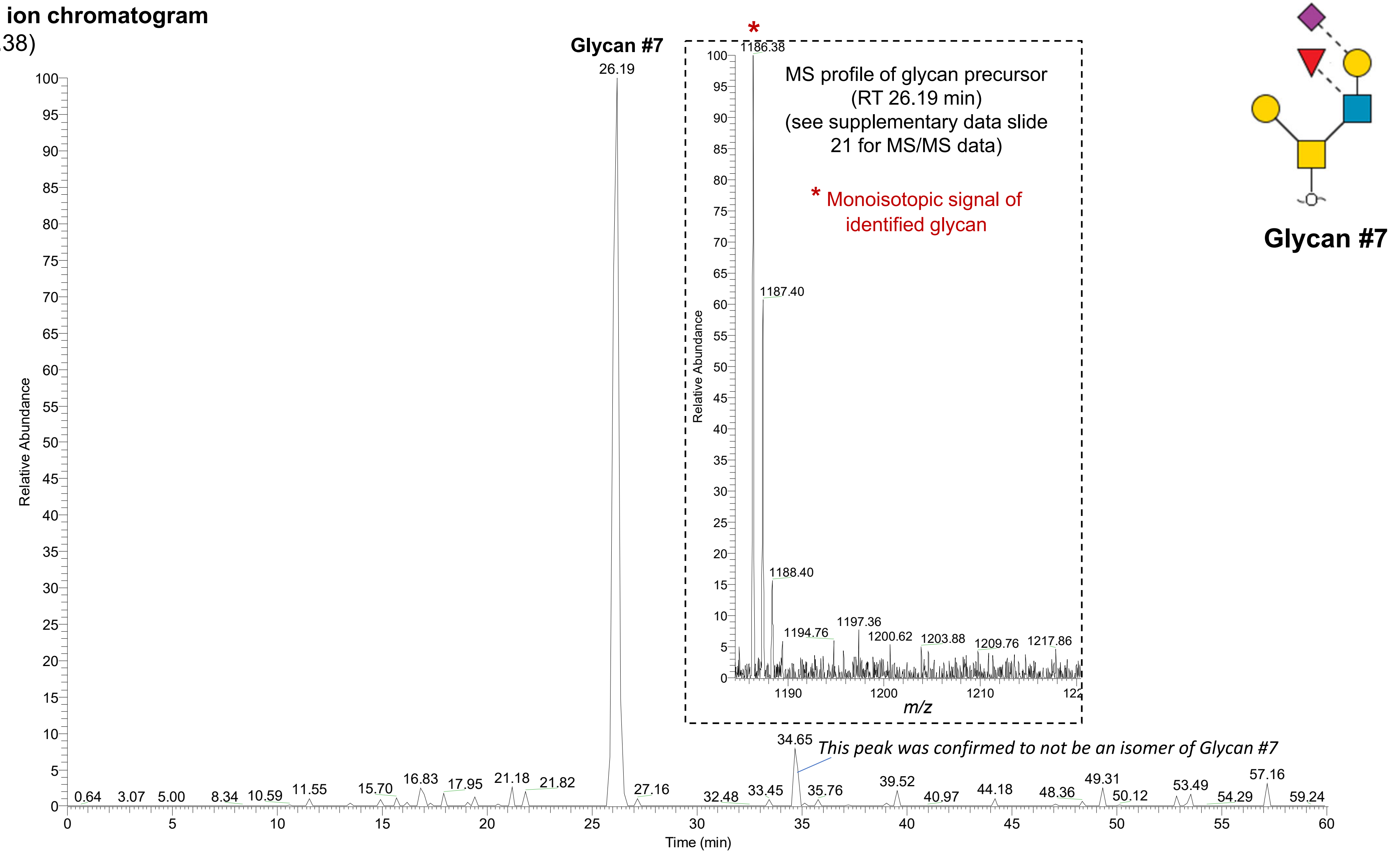

**Glycan #8**
**Extracted ion chromatogram**
(*m/z* 1257.40)

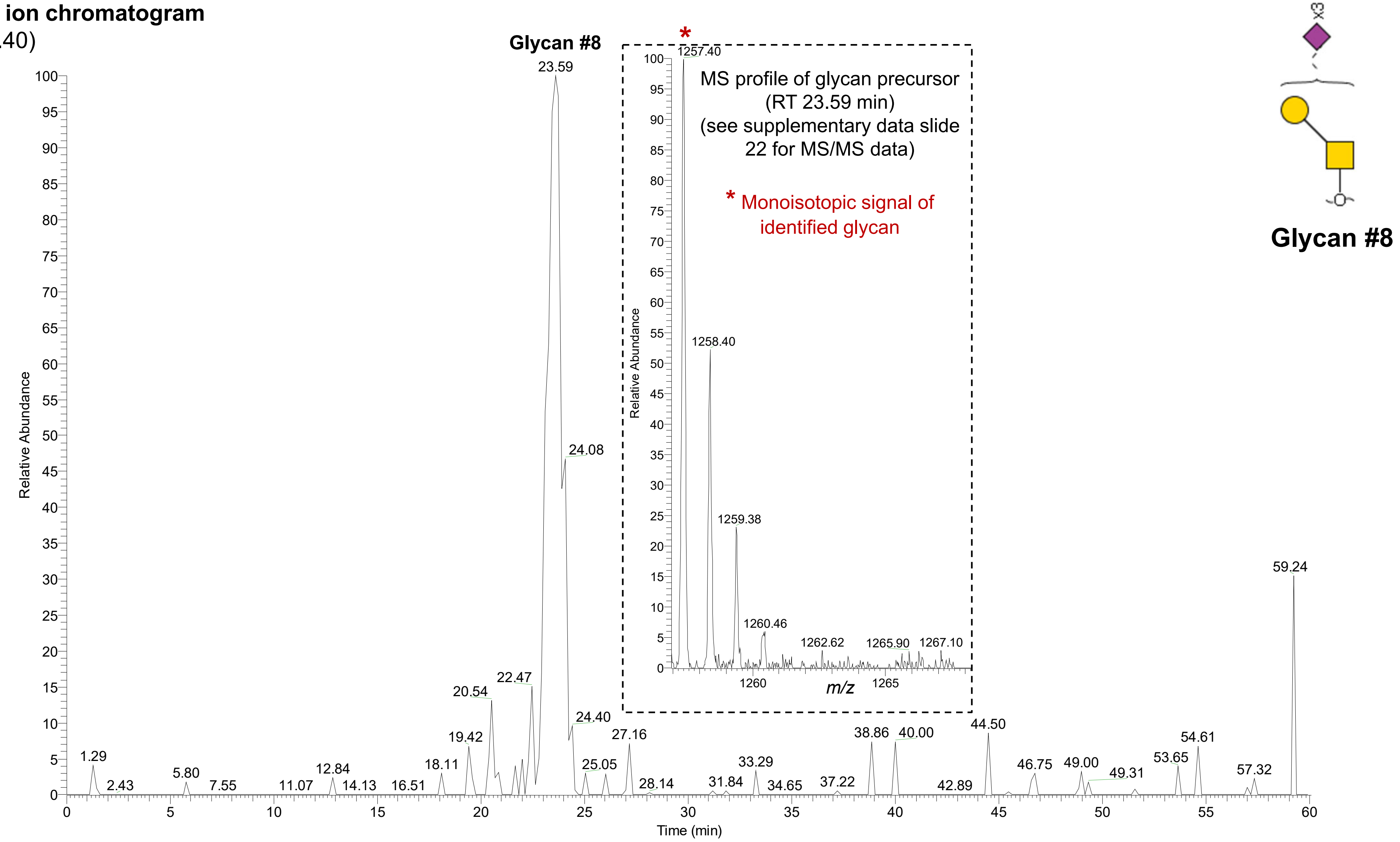

**Glycan #9**
**Extracted ion chromatogram**
(*m/z* 1331.44)

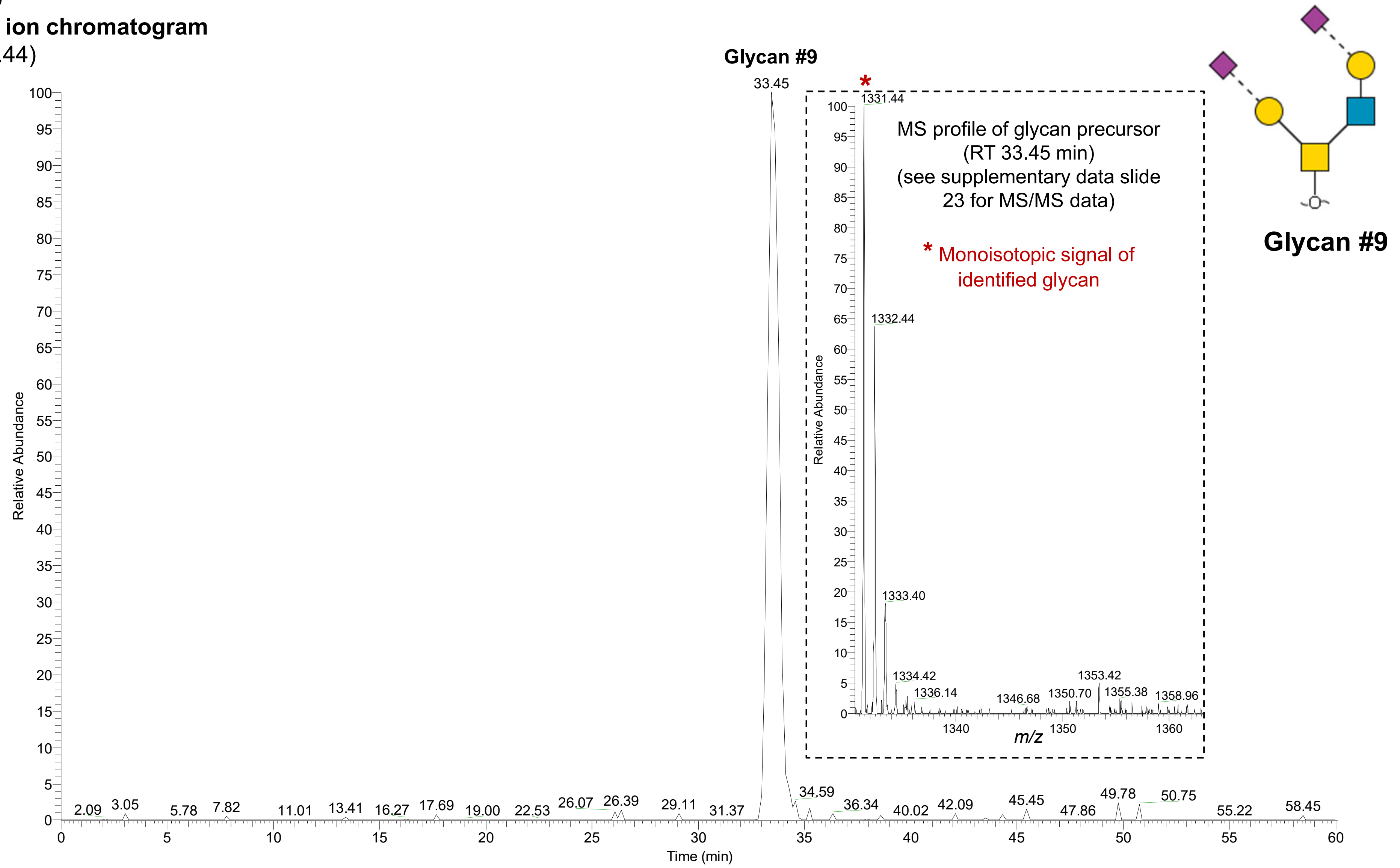

Manually annotated PGC-LC-ESI-CID-MS/MS (-)
spectra of *O*-glycans of platelet releasate activated with thrombin (0.2 U/mL)

**Glycan #1**
Observed *m/z* 327.12 (1-), RT: ~15.44 min
$[M-H]^-$ 327.12 Da

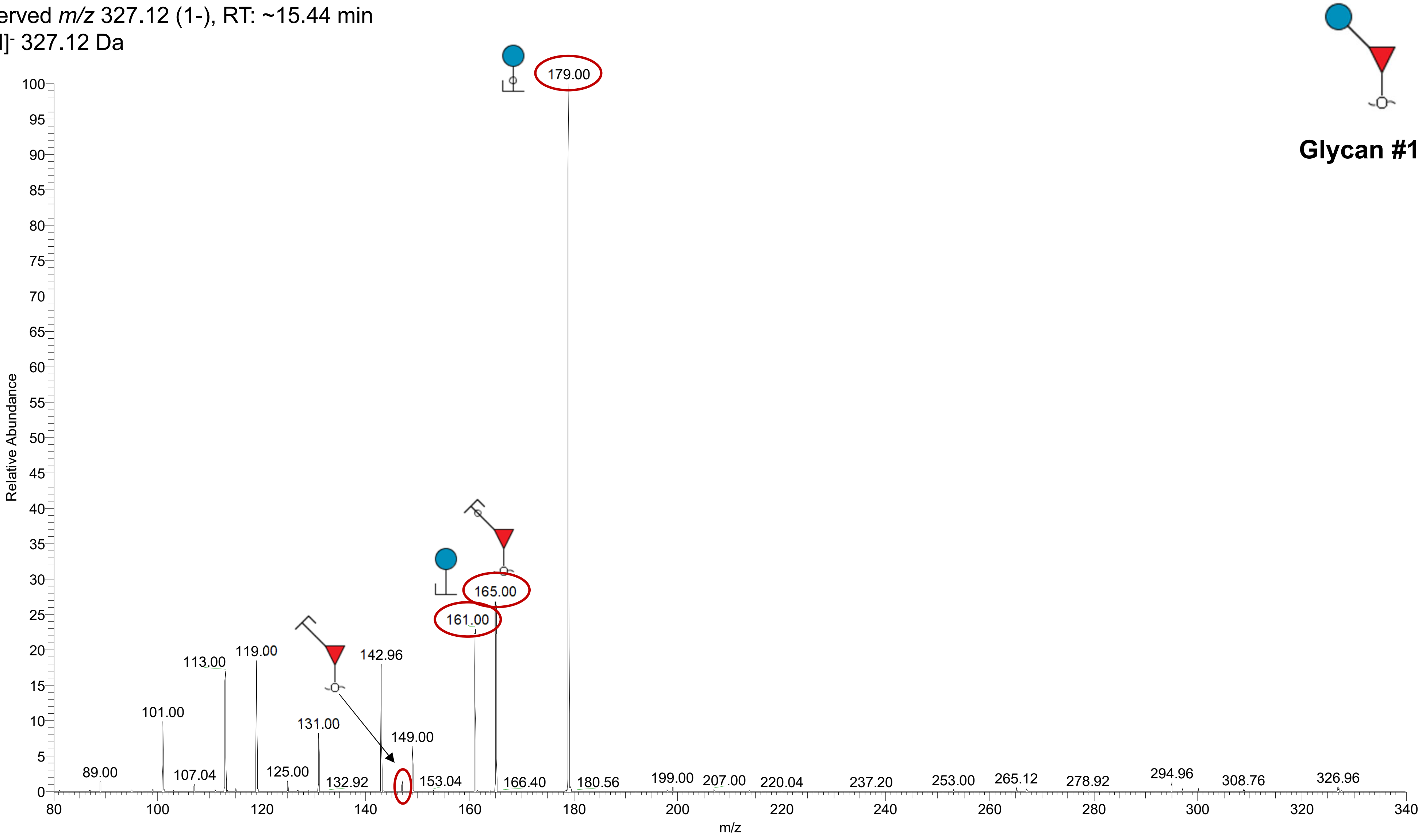

**Glycan #2**

Observed *m/z* 445.10 (1-), RT: ~15.68 min

$[M-H]^-$ 445.10 Da

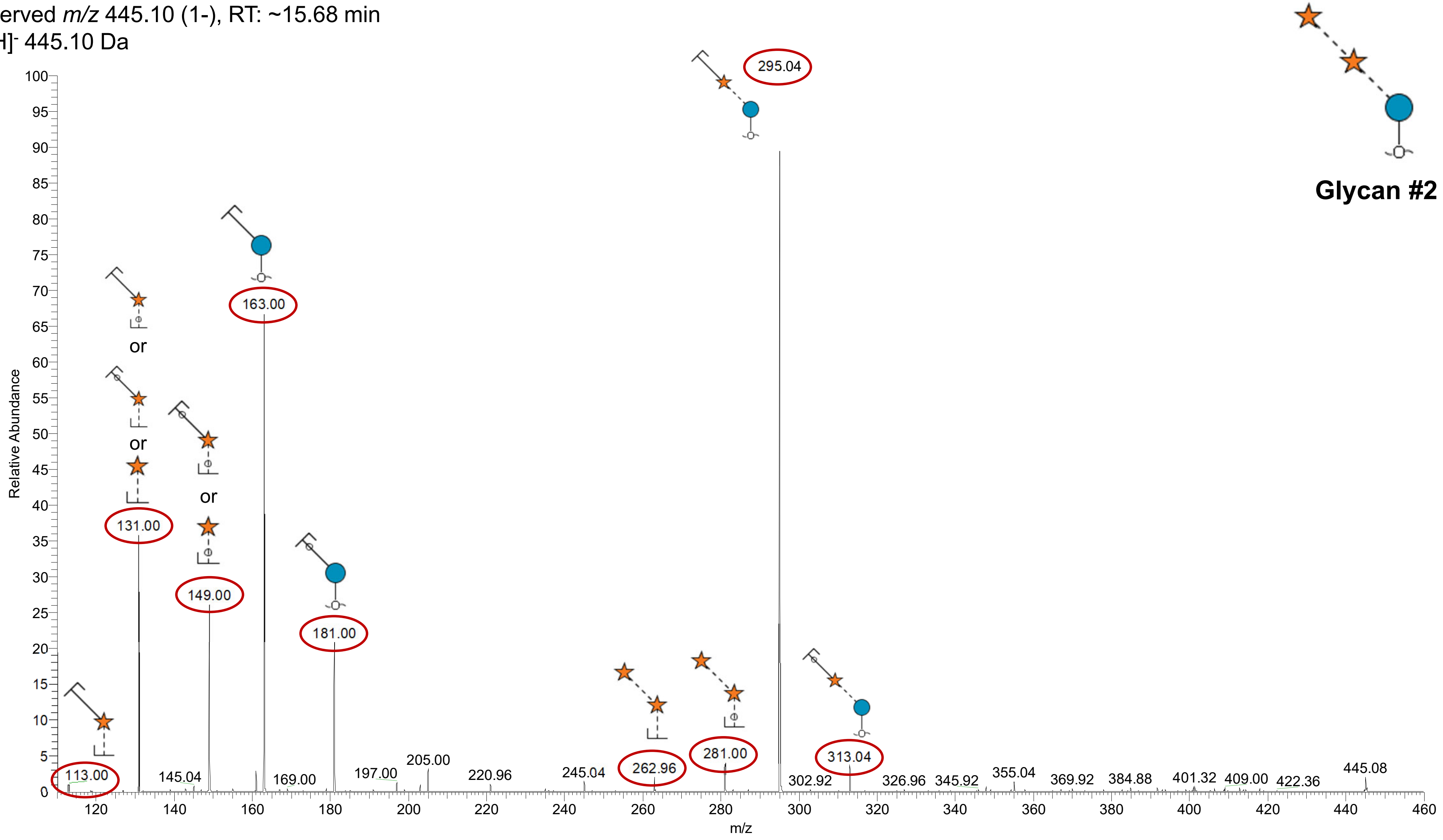

**Glycan #3a**
Observed *m/z* 675.16 (1-), RT: ~16.43 min
$[M\text{-}H]^-$ 675.16 Da

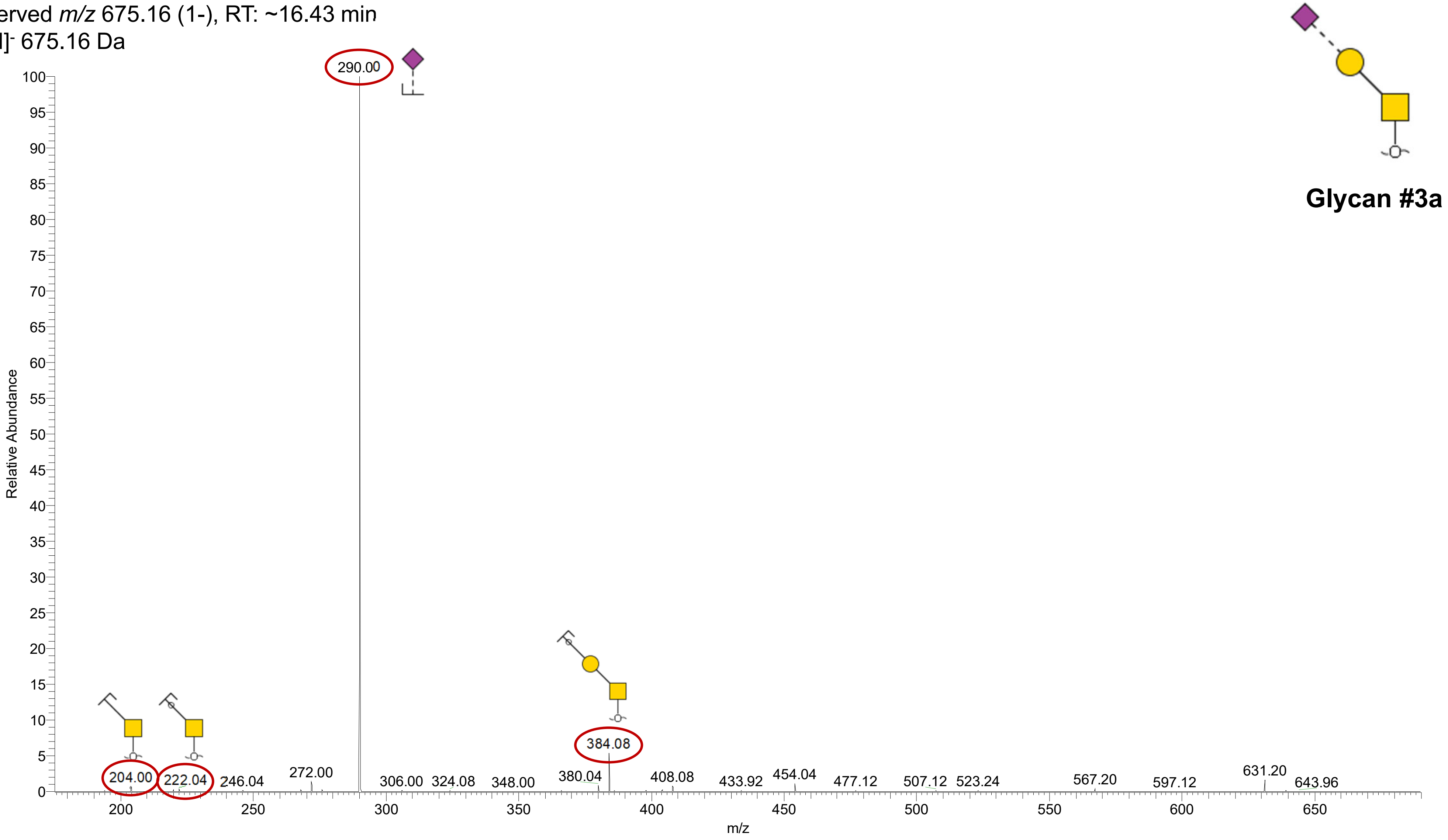

**Glycan #3b**
Observed *m/z* 675.16 (1-), RT: ~20.45 min
$[M\text{-}H]^-$ 675.16 Da

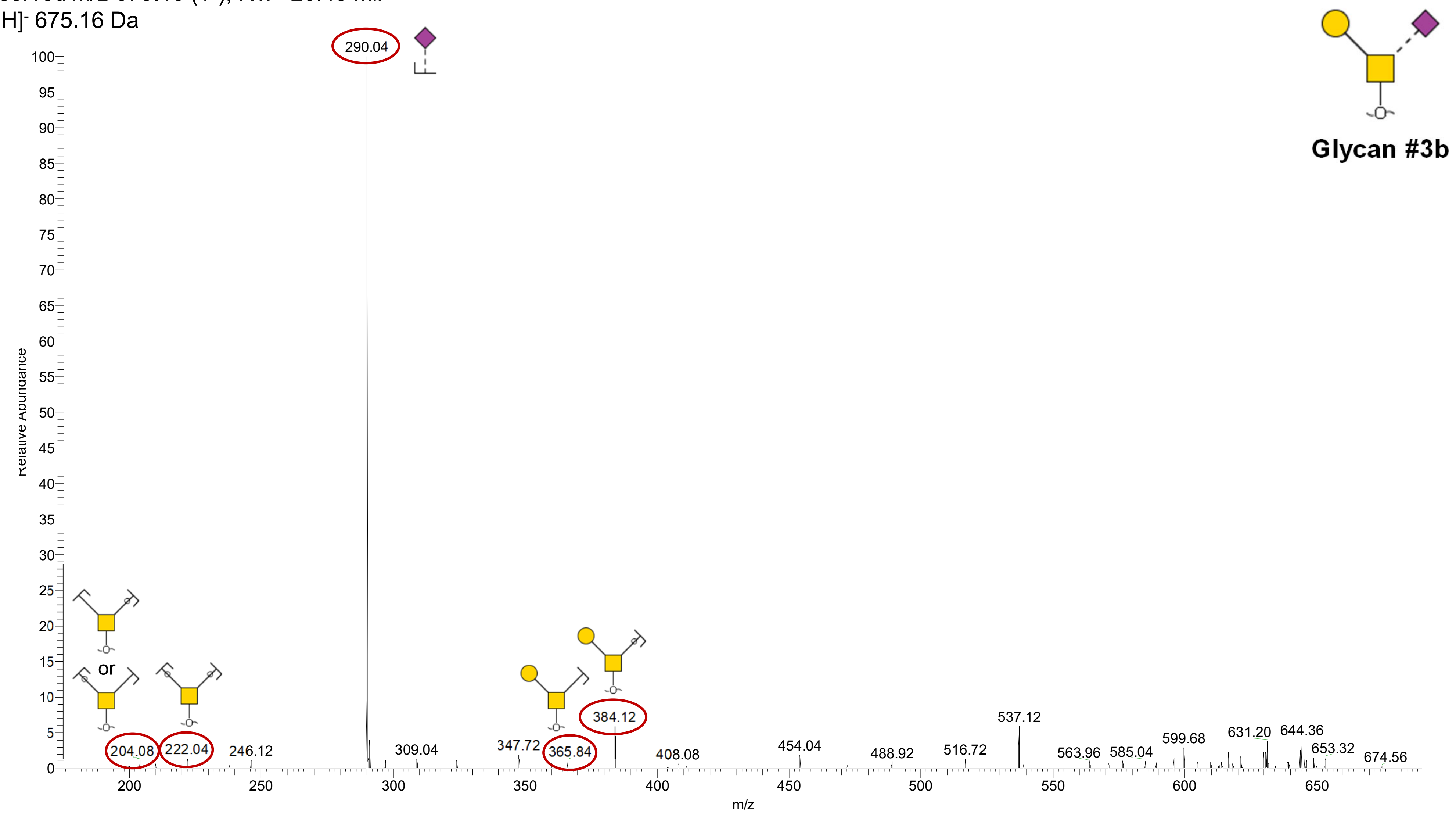

**Glycan #4**
Observed *m/z* 878.22 (1-), RT: ~18.49 min
$[M\text{-}H]^-$ 878.22 Da

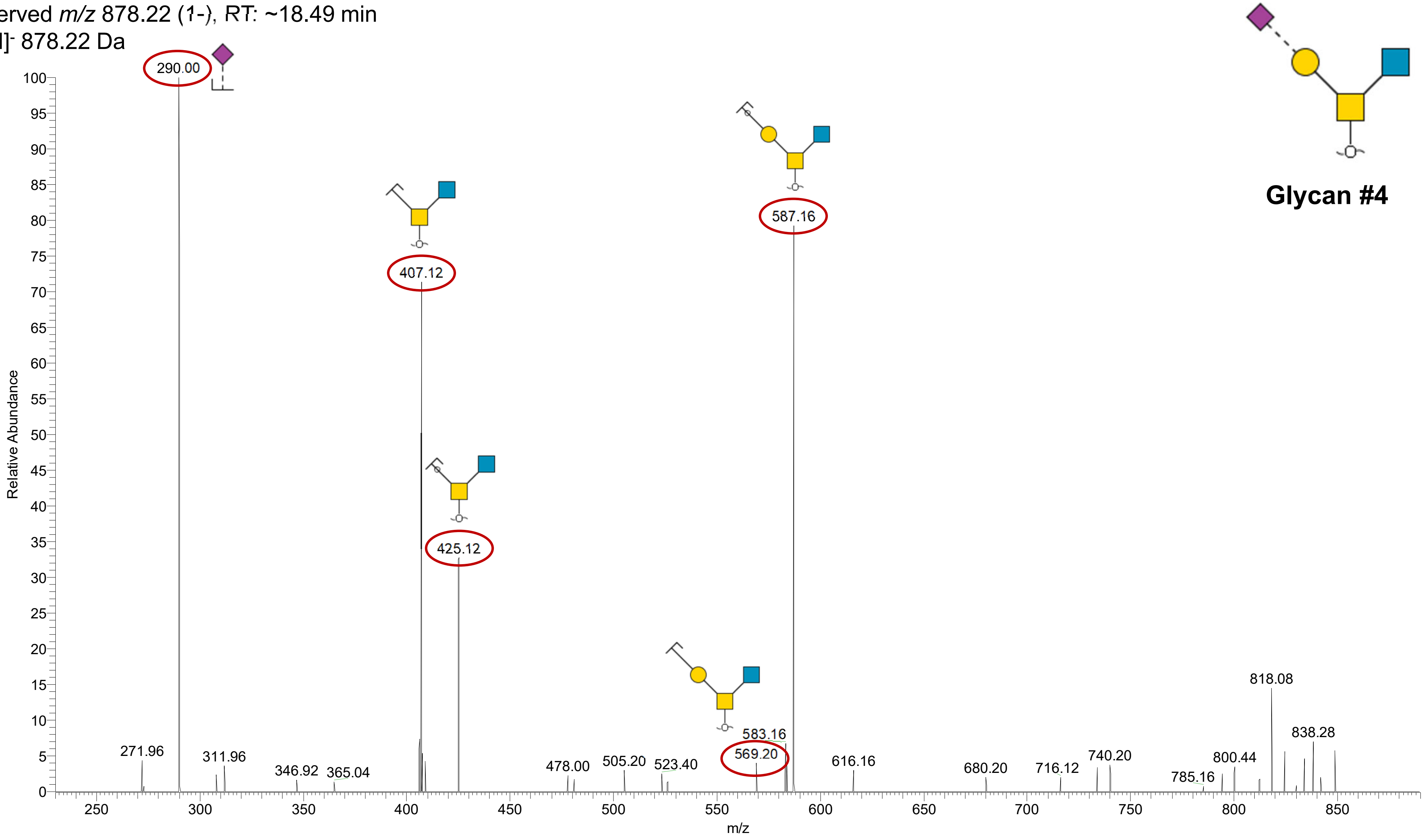

**Glycan #5**

Observed *m/z* 966.26 (1-), RT: ~18.35 min

$[M-H]^-$ 966.26 Da

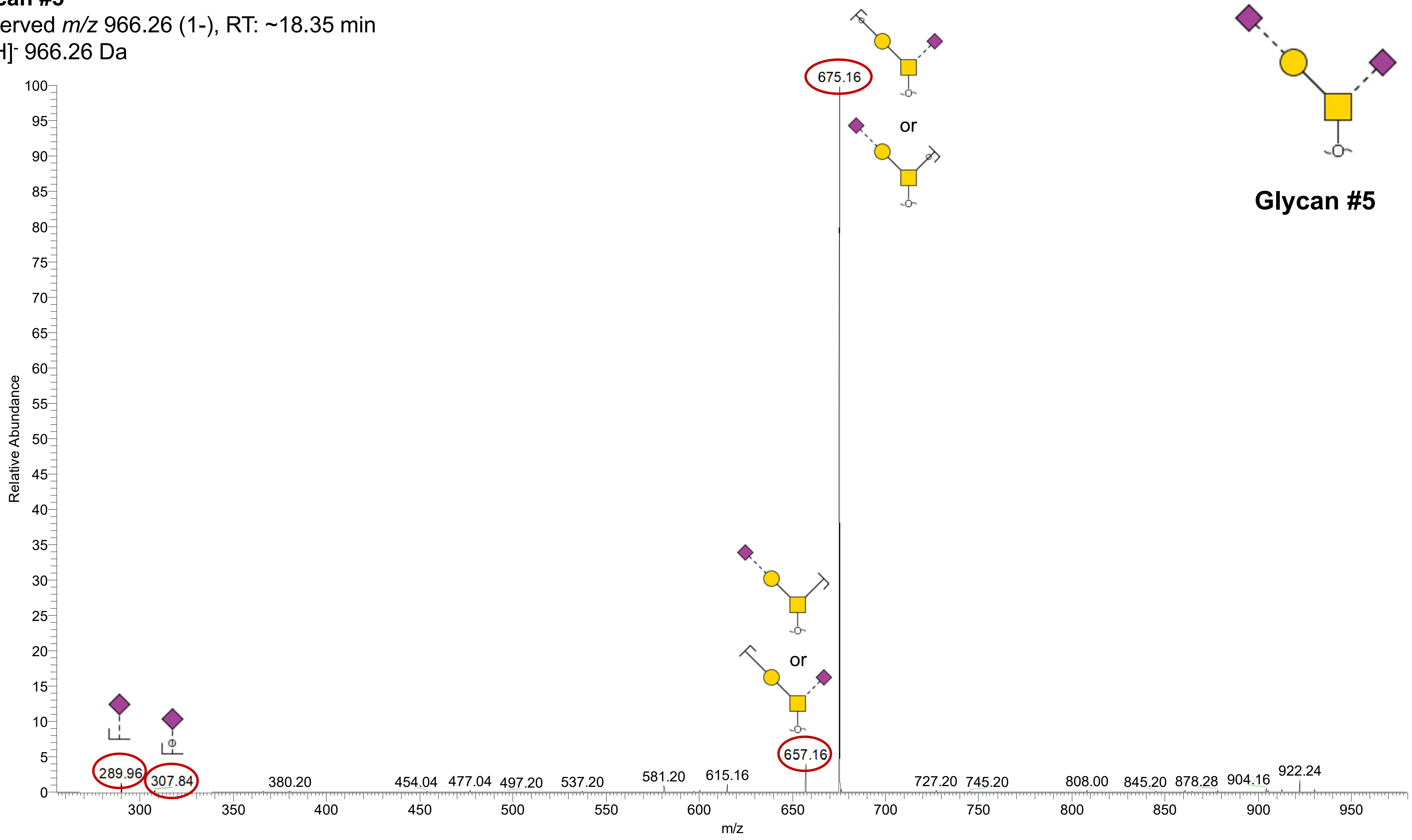

**Glycan #6a**

Observed *m/z* 1040.31 (1-), RT: ~21.39 min

$[M\text{-}H]^-$ 1040.32 Da

Note: The glycan isomers #6a and #6b are only distinguishable via PGC-LC retention time.

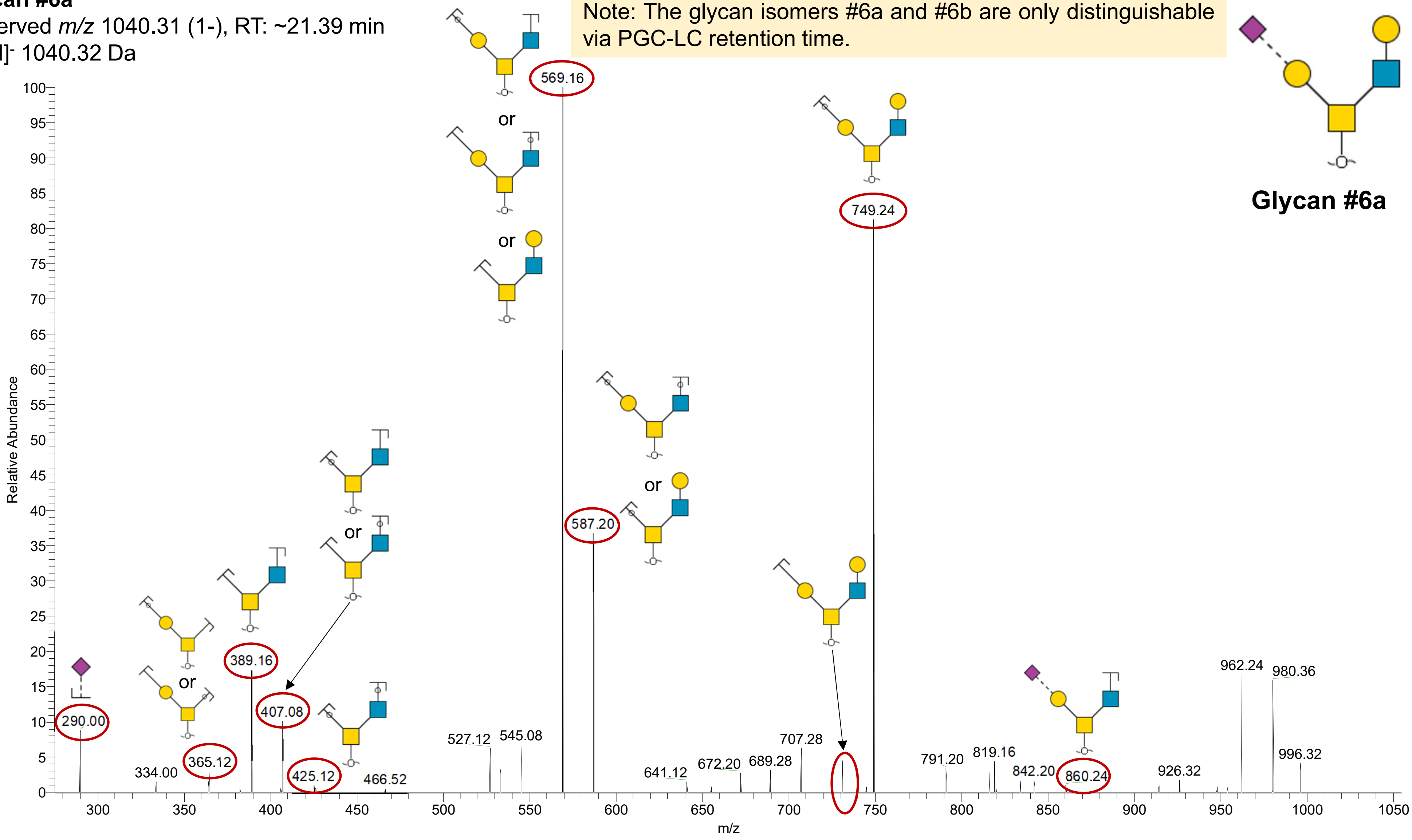

**Glycan #6b**
Observed *m/z* 1040.31 (1-), RT: ~26.16 min
[M-H]$^-$ 1040.32 Da

Note: The glycan isomers #6a and #6b are only distinguishable via PGC-LC retention time.

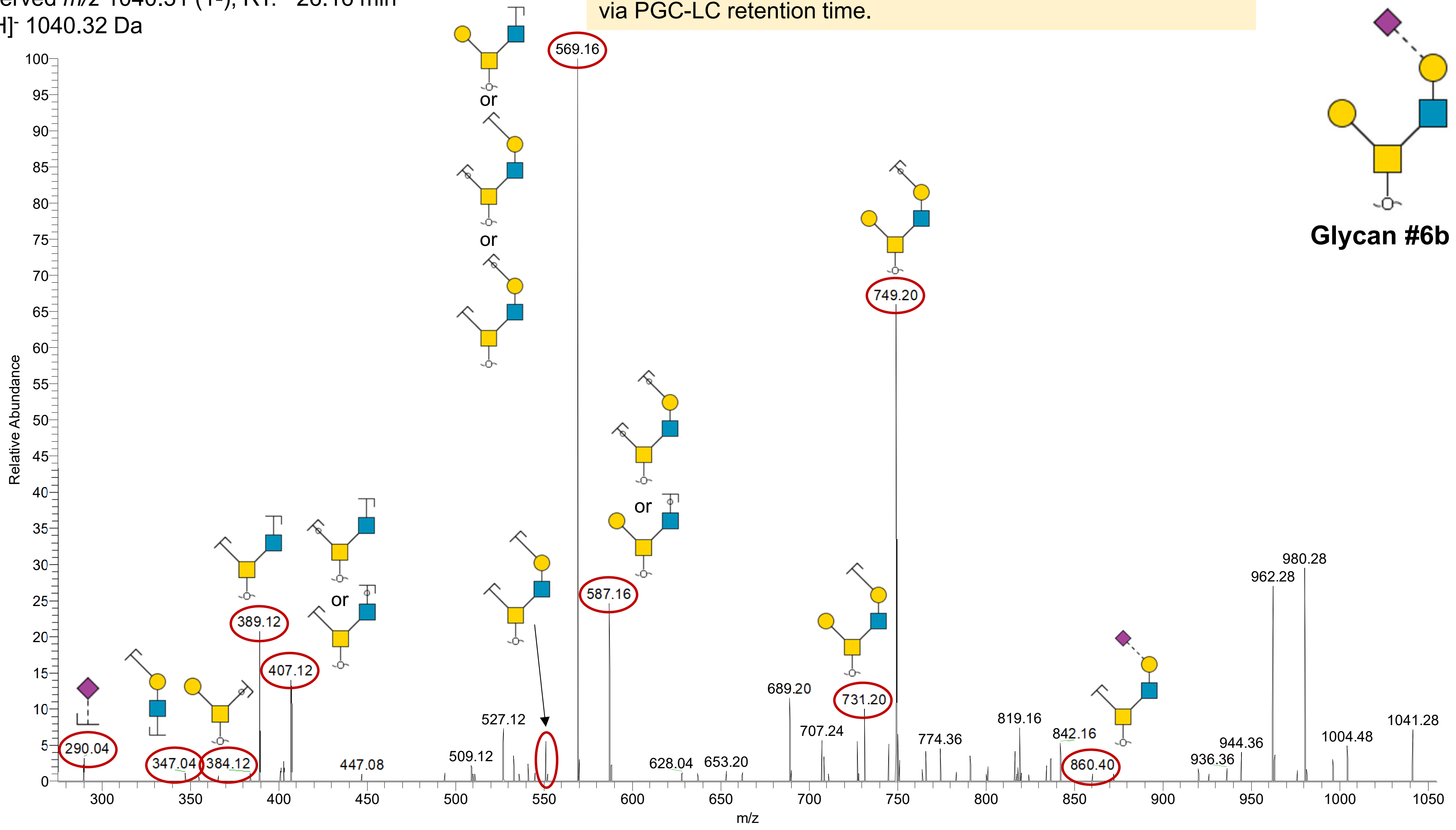

**Glycan #7**
Observed *m/z* 1186.38 (1-), RT: ~26.06 min
$[M\text{-}H]^-$ 1186.38 Da

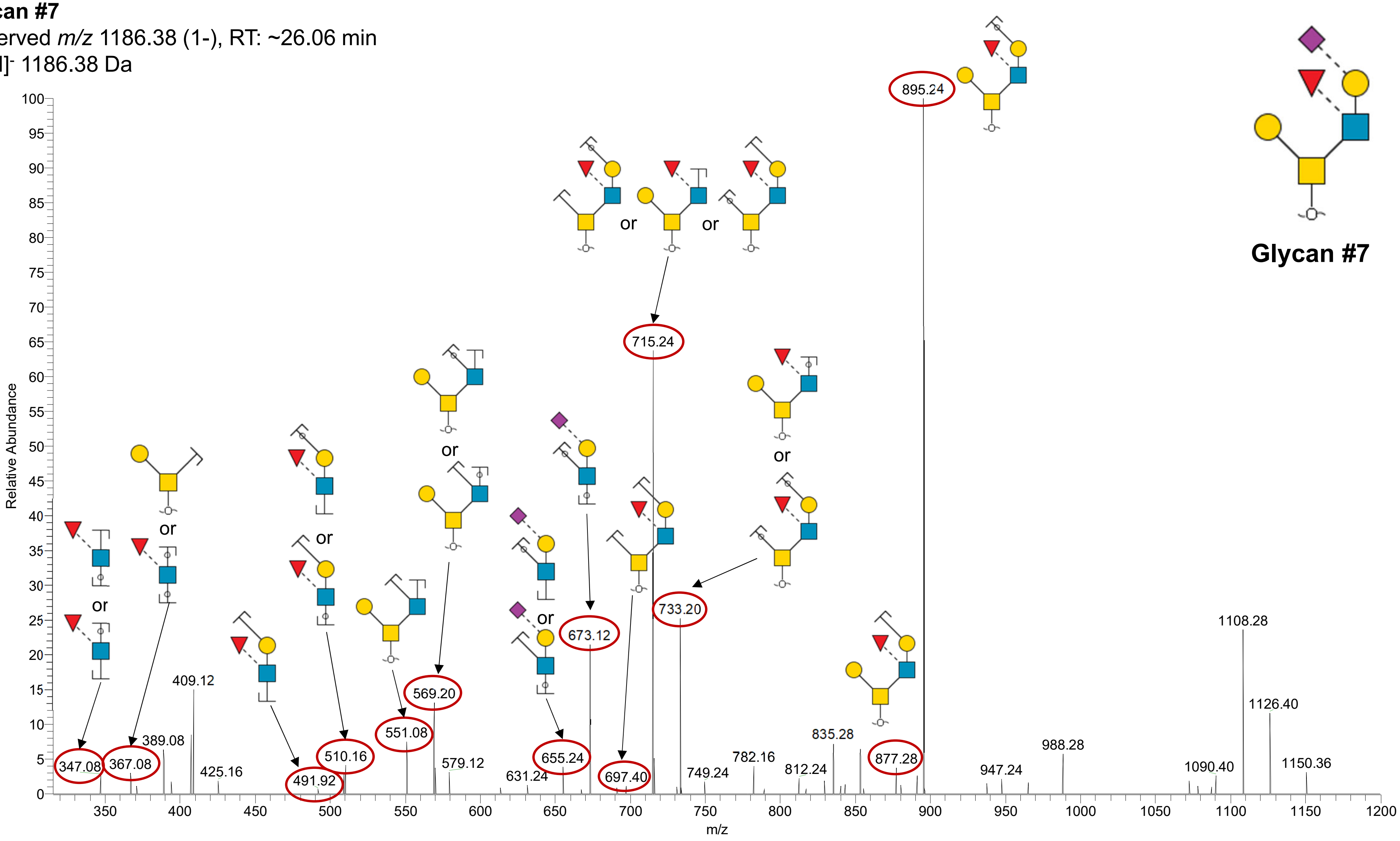

**Glycan #8**
Observed *m/z* 1257.40 (1-), RT: ~24.17 min
$[M-H]^-$ 1257.38 Da

Note: This glycan has not been annotated with sialyl linkages due to not enough supporting evidence from the PGC-LC elution pattern.

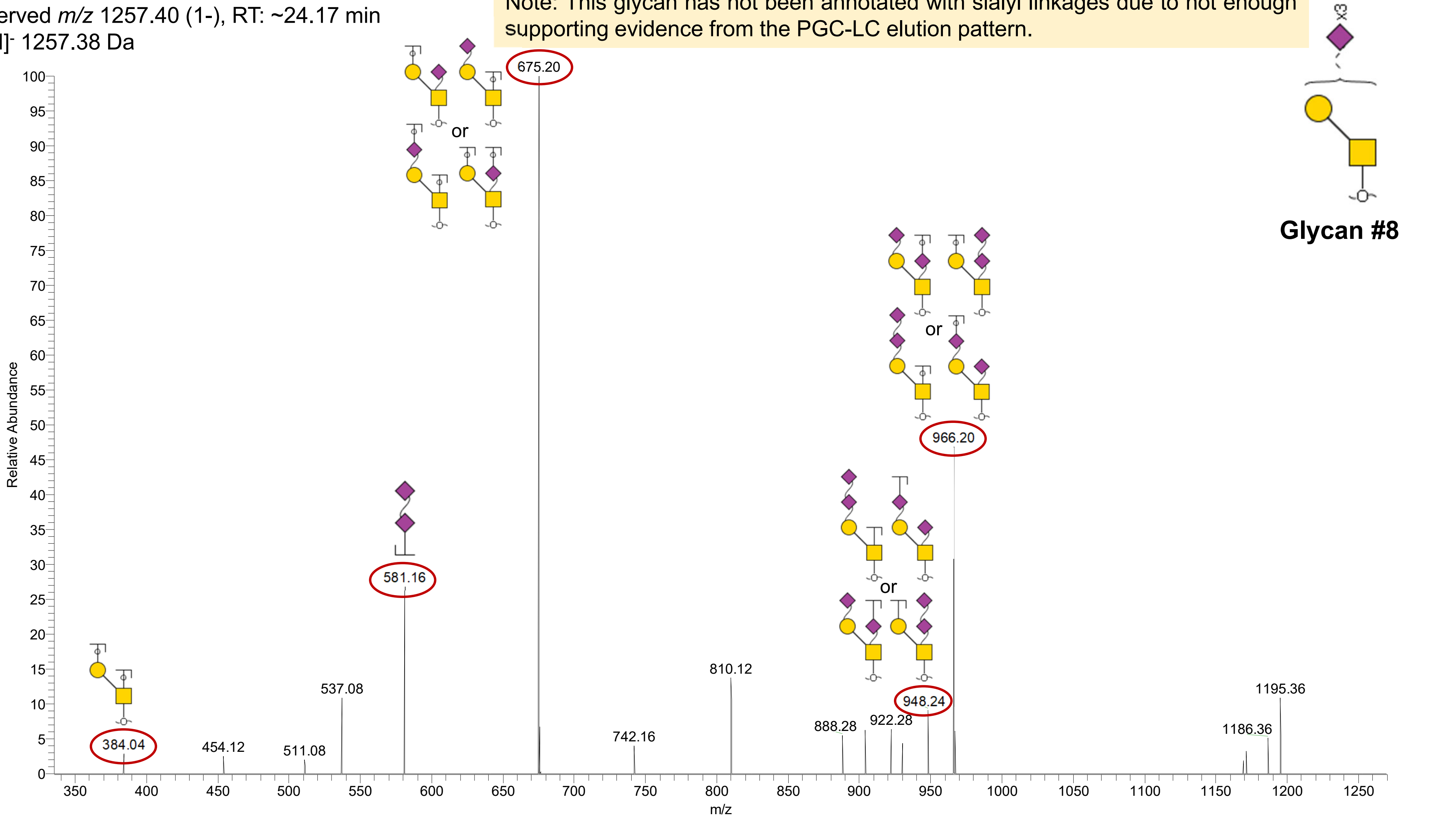

**Glycan #9**
Observed *m/z* 1331.44 (1-), RT: ~33.50 min
[M-H]⁻ 1331.42 Da

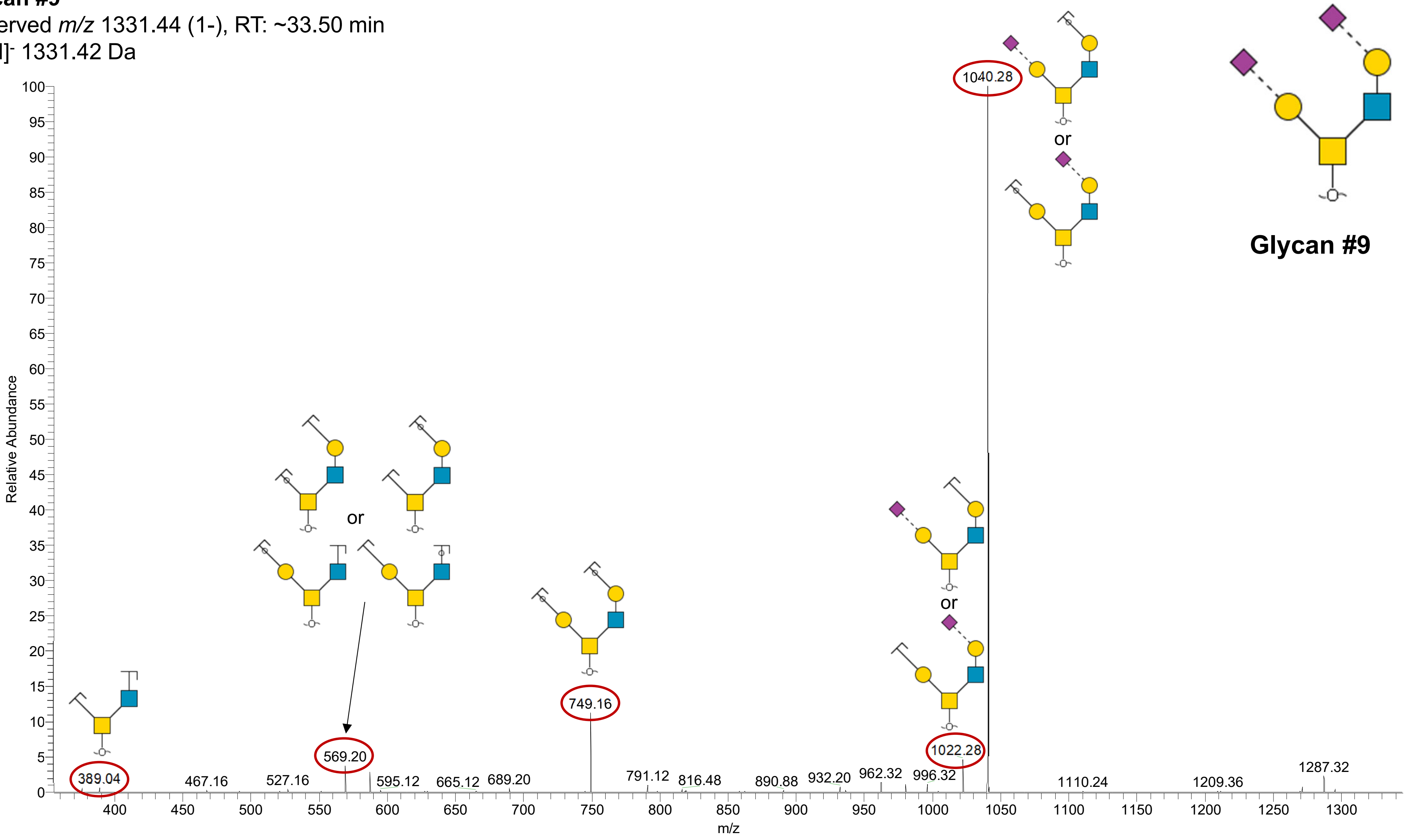
